# T follicular helper cell profiles differ by malaria antigen and for children compared to adults

**DOI:** 10.1101/2024.04.13.589352

**Authors:** Catherine S. Forconi, Christina Nixon, Hannah W. Wu, Boaz Odwar, Sunthorn Pond-Tor, John M. Ong’echa, Jonathan Kurtis, Ann M. Moormann

## Abstract

**Background:** Circulating T-follicular helper (cT_FH_) cells have the potential to provide an additional correlate of protection against *Plasmodium falciparum* (*Pf)* as they are essential to promote B-cell production of long-lasting antibodies. Assessing the specificity of cT_FH_ subsets to individual malaria antigens is vital to understanding the variation observed in antibody responses and identifying promising malaria vaccine candidates.

**Methods:** Using spectral flow cytometry and unbiased clustering analysis, we assessed antigen-specific cT_FH_ cell recall responses *in vitro* to malaria vaccine candidates *Pf-*schizont egress antigen-1 (*Pf*SEA-1A) and *Pf*-glutamic acid-rich protein (*Pf*GARP) within a cross-section of children and adults living in a malaria-holoendemic region of western Kenya.

**Findings:** In children, a broad array of cT_FH_ subsets (defined by cytokine and transcription factor expression) were reactive to both malaria antigens, *Pf*SEA-1A and *Pf*GARP, while adults had a narrow profile centering on cT_FH_17- and cT_FH_1/17-like subsets following stimulation with *Pf*GARP only.

**Interpretation:** Because T_FH_17 cells are involved in the maintenance of memory antibody responses within the context of parasitic infections, our results suggest that *Pf*GARP might generate longer-lived antibody responses compared to *Pf*SEA-1A. These findings have intriguing implications for evaluating malaria vaccine candidates as they highlight the importance of including cT_FH_ profiles when assessing interdependent correlates of protective immunity.

## Introduction

Despite progress made to reduce the global malaria burden, *Plasmodium falciparum* (*Pf*) remains one of the leading causes of mortality among children under 5 years of age.^1^ Unfortunately, progress has been impeded by a plateau in malaria control since 2015. Anti-malarial drug resistance,^2,3^ and unprecedented logistical challenges during the COVID-19 pandemic that dramatically impacted the distribution of insecticide-impregnated mosquito nets led to an increase in malaria cases in children under 5 years of age from 17.6 million in 2015 to 19.2 million in 2021 in East and Southern African countries,^1,4^ thus, contributing to reinvigorated prioritization of malaria vaccine initiatives.

After 30 years of development, the first malaria vaccine—RTS,S/AS01E—was approved by the World Health Organization in 2021 for use in children residing in malaria-endemic regions. However, the limited efficacy of RTS,S,^5^ particularly against severe malaria^5^, has motivated the search for additional vaccine candidate antigens, including blood-stage *Pf*-schizont egress antigen-1 (*Pf*SEA-1A)^6^ and *Pf*-glutamic acid-rich protein (*Pf*GARP).^7^ Antibodies against *Pf*SEA-1A correlate with significantly lower parasite densities in Kenyan adults and adolescents and substantially reduce schizont replication *in vitro*.^6,8^ Whereas *Pf*GARP-specific antibodies kill trophozoite-infected erythrocytes in culture and confer partial protection against *Pf*-challenge in vaccinated non-human primates.^7^ The presence of antibodies against *Pf*GARP was associated with a 2-fold-lower parasite density in Kenyan adults and adolescents.^7^ Due to the complexity of the parasites’ life cycle, there is consensus that a multi-valent vaccine would be more efficacious^9^. However, determining which vaccine candidate antigens hold the most promise for inclusion in next-generation malaria vaccines remains a challenge.

Many pathogens and vaccines engender protective antibody responses after a single or few exposures^10^, characterized by the production of long-lived plasma cells and memory B cells.^11^ Affinity maturation takes place in the germinal center (GC), where antigen-specific CD4^pos^ T-follicular helper (T_FH_) cells are required not only to provide cellular (CD40L) and molecular (IL-21, IL-4, and IL-13) signals to trigger B-cell proliferation but also to promote GC maintenance and plasmablast differentiation. The GC is also where higher affinity B cells outcompete B cells with lower affinity for T_FH_ help.^12,13^ T_FH_ cells are defined by the expression of a combination of markers, starting with chemokine receptor CXCR5, which directs CD4^pos^ T cells from the T-cell zone to engage with follicular B cells.^14,15^ Once antigen-experienced T_FH_ cells leave the GC, they become circulating T_FH_ (cT_FH_) cells and correlate with the generation of long-lasting antibody responses. Interestingly, cT_FH_ cells can also come from peripheral cT_FH_ precursor CCR7^low^PD1^high^CXCR5^pos^ cells; thus, they also have a GC-independent origin^16^. Expression profiles of CCR6 and CXCR3 categorize cT_FH_ subsets as follows: cT_FH_1 (CCR6^neg^CXCR3^pos^), cT_FH_2 (CCR6^neg^CXCR3^neg^), and cT_FH_17 (CCR6^pos^CXCR3^neg^)^17^, whereas the expression of PD-1, ICOS, CD127, and CCR7 define their functional status: quiescent/central memory cT_FH_ (CCR7^high^PD-1^neg^ICOS^neg^CD127^pos^) or activated/effector memory cT_FH_ (CCR7^low^PD-1^pos^ICOS^pos^CD127^low/neg^).^17–19^ In addition, transcription factors (i.e., Bcl6 and cMAF) and cytokines (i.e., interferon-gamma [IFN□], IL-4, and IL-21) are crucial to further characterizing the role of each cT_FH_ subset within the context of their interactions with B cells to promote antibody production.^20–23^ Bcl6 facilitates the production of IL-21 by T cells which aids B-cell affinity maturation and antibody production.^24–26^ T_FH_ cells also secrete cytokines that align with their subset classifications (but are not limited to them), such as IFN□ (T_FH_1), IL-4/IL-13 (T_FH_2), IL-4/IL-5/IL-13 (T_FH_13), and IL-17 (T_FH_17), that direct antibody isotype class-switching and mediate effector functions.^12,18,20–22,27^

Several studies have characterized cT_FH_ subsets within the context of both adult and childhood *Pf*-malaria infections, as well as in healthy malaria-naïve volunteers under controlled malaria infection conditions.^28^ In Mali, where malaria is seasonal, Obeng-Adjei and colleagues showed that T_H_1-polarized cT_FH_ PD1^pos^CXCR5^pos^CXCR3^pos^ cells were preferentially activated in children and less efficient than CXCR3^neg^ cT_FH_ cells in helping autologous B cells produce antibodies; yet they used U.S. healthy adult cT_FH_ cells for their *in vitro* assays. Therefore, the authors suggested that promoting T_H_2-like CXCR3^neg^ cT_FH_ subsets could improve antimalarial vaccine efficacy.^29^ Similarly, a recent study in Papua, Indonesia, where malaria is perennial, found that all cT_FH_ subsets from adults showed higher activation and proliferation compared to cT_FH_ cells from children.^30^ After *in vitro* stimulation with infected red blood cells, they found that all cT_FH_ subsets were activated in adults accompanied by IL-4 production, whereas only the T_H_1-polarized cT_FH_ subset responded in children.^30^ One study in Uganda, where malaria is holoendemic with seasonal peaks, showed that a shift from T_H_2- to T_H_1-polarized cT_FH_ subsets occurred during the first 6 years of life and was associated with the development of functional antibodies against *Pf*-malaria, yet appeared to be independent of malaria exposure as this age-associated shift was also observed in a malaria-naïve population.^31^ Interestingly, this study also found that a higher proportion of T_H_17-cT_FH_ cells was associated with a decreased risk of *P. falciparum* infection the following year; however, the authors postulated that this phenomenon could have been driven by previous exposure to the parasite. The observed higher abundance of T_H_2-like cT_FH_ cells in children younger than 6 years old and the age-associated increase in T_H_1-cT_FH_ cells achieving the same proportion of T_H_1-cT_FH_ cells by 6 years of age and into adulthood appeared to be independent of malaria exposure. Of note, these studies were limited by the use of a 2-dimensional gating strategy to classify cT_FH_ subsets and whole parasite activation conditions, either during infections or *in vitro* stimulation assays, leaving the malaria-antigen specificity of the different cT_FH_ subsets responses undefined. However, these studies highlight the progression in the field of evaluating the role of human T_FH_ cells in anti-malarial immunity.

Antibody levels are an unreliable predictor of malaria vaccine efficacy.^32^ Thus, establishing a combined immune profile to incorporate other surrogates of protection is warranted. Here, we examined the profile of cT_FH_ subsets using multiparameter spectral flow cytometry^33,34^ against two malaria vaccine candidate antigens (*Pf*SEA-1A^6^ and *Pf*GARP^7^) in a cross-section of children and adults residing in a malaria-holoendemic region of Kenya. Our findings revealed significant differences in cT_FH_ subsets between children and adults, where children had more abundant cT_FH_1-like cells and showed antigen-specific responses from all cT_FH_ types, whereas these responses were limited to cT_FH_17- and 1/17-like cells in adults. Moreover, this study showed that *Pf*GARP triggered Bcl6 expression across cT_FH_ subsets, whereas *Pf*SEA-1A induced more cMAF expression, demonstrating the feasibility of implementing T-cell immune correlates to down-select new malaria candidates.

## Results

### Children had lower anti-PfSEA-1A antibodies compared to adults but similar levels of anti-PfGARP antibodies

This cross-sectional study selected a convenience sample of 7-year-old children and adults with a mean age of 22.67 years (ranging from 19 to 30 years old). No statistical difference was observed (*p*=0.35 after Welch’s test) regarding the absolute lymphocyte count (ALC) between children (mean of 373.1; standard deviation [SD] of 118.1) and adults (mean of 335.2; SD of 99.08). Children had significantly lower hematocrit values (median of 37%, interquartile range [IQR] of 32.7–40.5) compared to adults (median of 44.4%, IQR of 40.5– 46), *p=*0.002 after a two-tailed Mann-Whitney *t*-test; however, these values were within normal ranges after adjusting for age.^35^ Both males and females (sex assigned at birth) were enrolled, with 43% (6/14) and 53% (8/15) being female within the children and adult groups, respectively. Seroprofiles against a panel of commonly used malaria antigens were generated to confirm the history of previous malaria infections for the selected children (Figure 1a). All participants had high IgG levels against merozoite antigens, apical membrane antigen 1 (AMA-1) and merozoite surface protein (MSP1), confirming at least one malaria infection within their lifetime.^36,37^ IgG antibodies against circumsporozoite protein (CSP) and CelTOS (liver-stage antigens) were characteristically lower than against blood-stage antigens yet were present in all study children. We observed a clear bimodal distribution in antibody levels against histidine-rich protein 2 (HRP2) with half of the children having “high-HRP2” versus “low-HRP2” IgG levels, possibly a reflection of recent malaria history since it has been suggested that HRP2-specific antibodies are short-lived and could serve as a surrogate for a recent infection.^38^ We assessed serological profiles for two *Pf*-malaria antigens being considered as potential vaccine candidates,^6,7^ *Pf*SEA-1A and *Pf*GARP (Figure 1b). We found that children had a significantly lower median level of IgG against *Pf*SEA-1A compared to adults (*p*<0.0001), whereas median levels to *Pf*GARP were similarly high for adults and children, yet with a broad range of reactivity.

**Figure 1:**
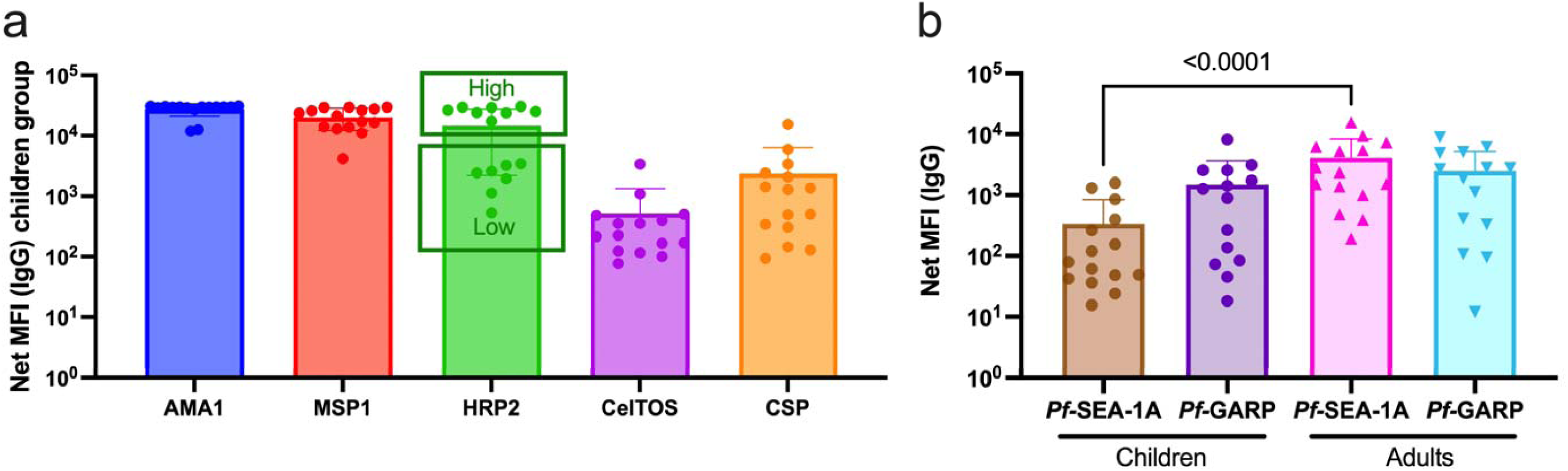
*Pf-*malaria IgG sero-profiles for children and adults. **(a)** IgG antibody levels against AMA1, MSP1, HRP2, CelTos, and CSP for children (n=14). The bar plots indicate the mean with SD. (**b)** IgG antibody levels against *Pf*SEA-1A and *Pf*GARP comparing children (n=15) and adults (n=15). The bar plots indicate the mean with SD. Net MFI values are the antigen-specific MFI values minus the BSA background. Mann-Whitney tests were performed.

### The overall abundance of CD4^pos^CXCR5^pos^ cells is unaltered by in vitro antigen stimulation

To determine whether *in vitro* antigen stimulation with *Pf*SEA-1A or *Pf*GARP altered the abundance of total cT_FH_ cells, we compared cT_FH_ cells from adults and children using a FlowSOM unbiased clustering analysis and EMBEDSOM dimensional reduction based on common lineage markers assessed by spectral flow cytometry from 87,812 live lymphocytes from each sample: CD8^pos^, CD4^pos^, CD4^pos^CXCR5^pos^, and CD4^pos^CD25^pos^ (Figure 2a). As expected, after a short stimulation (6 hrs), overall abundances of cT_FH_ cells were similar across conditions for both adults (Figure 2b) and children (Figure 2c). This observation was confirmed by EdgeR statistical analysis (Supplemental Figure 2) and demonstrated that *in vitro* stimulation did not preferentially expand the T cell populations on which we based our subsequent analyses.

**Figure 2:**
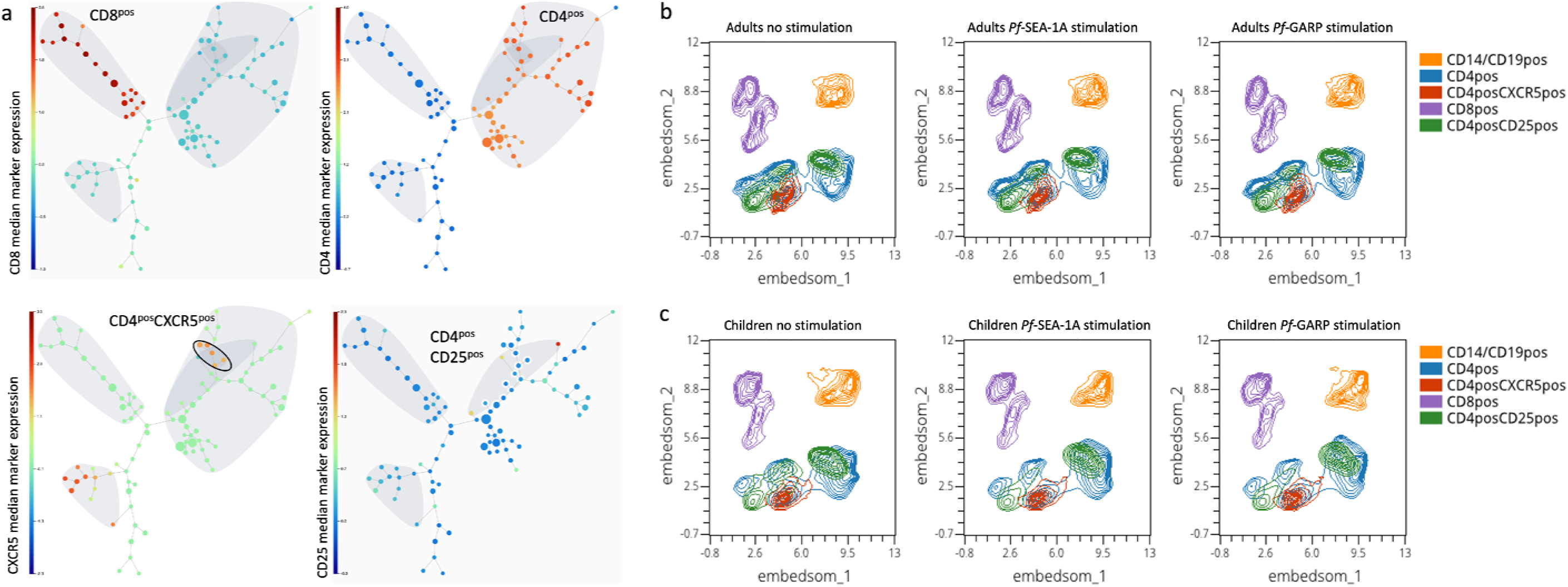
Cluster visualization of immune cell types and their abundance. **(a)** A 100- node and 75-meta-cluster FlowSOM tree was generated on live lymphocytes from all our participants (n=29 children and adults), highlighting CD3^pos^CD8^pos^, CD3^pos^CD4^pos^, CD4^pos^CXCR5^pos^, and CD4^pos^CD25^pos^ cells. The 5 nodes of the CD4^pos^CXCR5^pos^CD25^neg^ population are circled and were used for the downstream analysis. The colored scale was based on the median arcsinh-transformed marker expression. UMAP plots showing 5 clusters defined as follows: CD14^pos^ and CD19^pos^ (orange), CD4^pos^ (blue), CD4^pos^CXCR5^pos^ (red), CD8^pos^ (purple), and CD4^pos^CD25^pos^ (green) from PBMCs isolated from **(b)** adults (n=15) and **(c)** children (n=14) that were unstimulated, stimulated with *Pf*SEA-1A, or stimulated with *Pf*GARP, respectively, from left to right.

### An unbiased clustering analysis identifies twelve distinct cT_FH_ meta-clusters

Numerous markers and a two-dimensional gating strategy have previously been used to determine the frequency of cT_FH_ subsets.^29–31^ To simultaneously account for the expression of 17 markers required to define cT_FH_ subsets, we used FlowSOM unbiased clustering analysis to determine the frequency of cT_FH_ subsets from a pool of 1,000 CD3^pos^CD4^pos^CXCR5^pos^CD25^neg^ cells from each sample (total of 13,000 CD3^pos^CD4^pos^CXCR5^pos^CD25^neg^ cells in both children and adults). Based on CXCR3 and CCR6 expression as well as the expression of effector/memory/activation markers (CCR7, CD127, PD1, and ICOS), cytokines (IFN□, IL-4, and IL-21), and transcription factors (Bcl6 and cMAF), we initially identified 15 meta-clusters within the CD4^pos^CXCR5^pos^ T cells (Figure 3a). First, we found different CXCR5 expression levels between meta-clusters (Figure 3b); CXCR5 is essential for cT_FH_ cells to migrate to the lymph nodes and interact with B-cells. Data presented in Figure 3 are from children; however, we found similar observations in the adult group (Supplemental Figure 3). Because CD45RA^pos^CXCR5^pos^ cells are likely naïve cells with transient low expression of CXCR5 yet high expression of CD45RA, we excluded 3 meta-clusters using these criteria (i.e., MC12, MC14, and MC15) (Figure 3c). Then, using the overall expression of CXCR3 and CCR6 across the cT_FH_ subsets (Figure 3d and 3e), we identified the remaining 12 clusters as follows: MC01 and MC02 were cT_FH_2-like; MC06 and MC07 were cT_FH_1-like; MC09 and MC11 were cT_FH_1/17-like; MC10 and MC13 were cT_FH_17-like. However, CXCR3 expression was not clearly delineated for some subsets and did not align with the conventional CCR6 vs. CXCR3 cytoplot (Figure 3f and 3g, Supplemental Figure 4). Based on the heatmap (Figure 3f), MC03, MC04, MC05, and MC08 clusters appear closer to cT_FH_2-like MC01 and MC02 clusters than cT_FH_1-like clusters MC06 and MC07, suggesting that they might be part of the cT_FH_2-like subset. But, based on their intensity of CXCR3 expression and their distribution across the CCR6 vs. CXCR3 cytoplot (Supplemental Figure 4), we defined MC03, MC04, MC05, and MC08 clusters as “undetermined” and would require additional cytokine and transcription factor analyses to fully categorize them.

**Figure 3:**
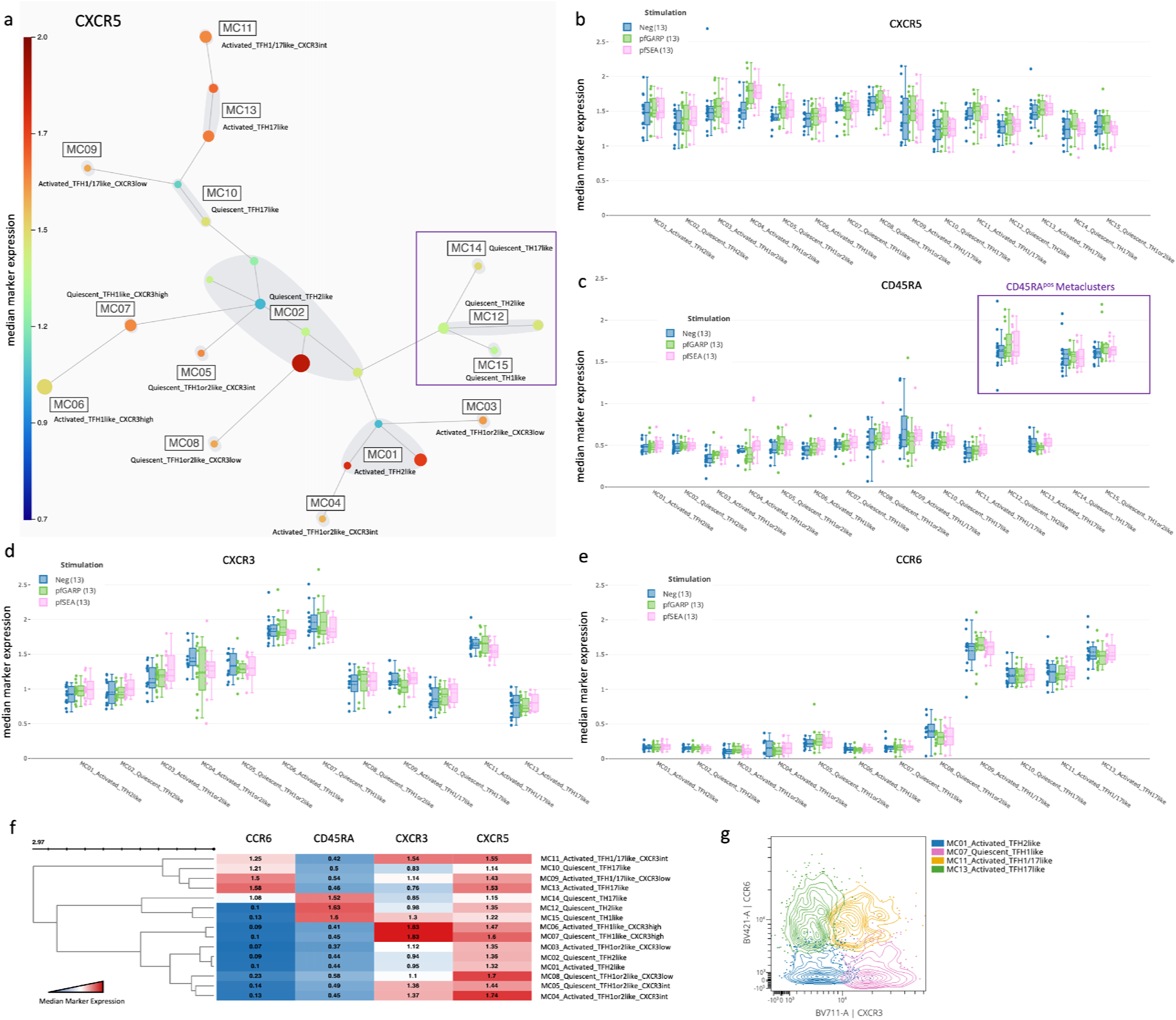
Characterization of T_FH_ subsets by CXCR5, CXCR3, CCR6, and CD45RA expression. **(a)** A new 25-node and 15 meta-cluster FlowSOM tree was generated from the 5 CD4^pos^CXCR5^pos^CD25^neg^ nodes shown in Figure 2A (CXCR5 expression tree). The colored scale of CXCR5 expression was based on the median arcsinh-transformed with red as the highest and deep blue as the absence of CXCR5 expression. **(b)** Box plots showing CXCR5 expression across all cT_FH_-like meta-clusters from children (n=13) with no stimulation (blue), and after *in vitro* stimulation with *Pf*GARP (green) or *Pf*SEA-1A (pink). Fifty percent of the data points are within the box limits, the solid line indicates the median, the dashed line indicates the mean, and the whiskers indicate the range of the remaining data with outliers being outside that range. Similar box plots are shown for **(c)** CD45RA, **(d)** CXCR3, and **(e)** CCR6 expression across all cT_FH_-like meta-clusters. **(f)** Clustered heatmap showing the median arcsinh-transformed expression for CCR6, CD45RA, CXCR3, and CXCR5 across meta-clusters, red showing the highest expression and blue the lowest. **(g)** Cytoplots of CCR6 vs. CXCR3 expression where MC01 is blue, MC07 is pink, MC11 is yellow, and MC13 is green.

### Heterogeneity of activation/maturation markers within cT_FH_ subsets

By assessing the expression of CCR7, PD1, CD127, and ICOS (Figure 4) and following the three-dimensional expression patterns adapted from Schmitt et al.^17^ (Supplemental Figure 5), we determined the activation state of each cT_FH_ meta-cluster and which markers created novel subsets. Data in Figure 4 are from children; however, similar observations were made for adults (Supplemental Figure 6). As expected, none of the extracellular markers showed significant differences in expression patterns after a short 6-hour stimulation, thus, representing the cT_FH_ repertoire present within our study participants. Interestingly, the cT_FH_1or2 subset (MC03) was the only meta-cluster with high expression of PD1 (Figure 4a) accompanied by high ICOS (Figure 4b), low CCR7 (Figure 4c), and low CD127 expression (Figure 4d), suggesting that MC03 was an activated/effector cT_FH_1or2 subset. The MC04 subset had low CCR7 and high ICOS expression but low CD127 and intermediate PD1 expression, indicating that this cluster was a less activated/effector cT_FH_1or2 cluster compared to MC03.

**Figure 4:**
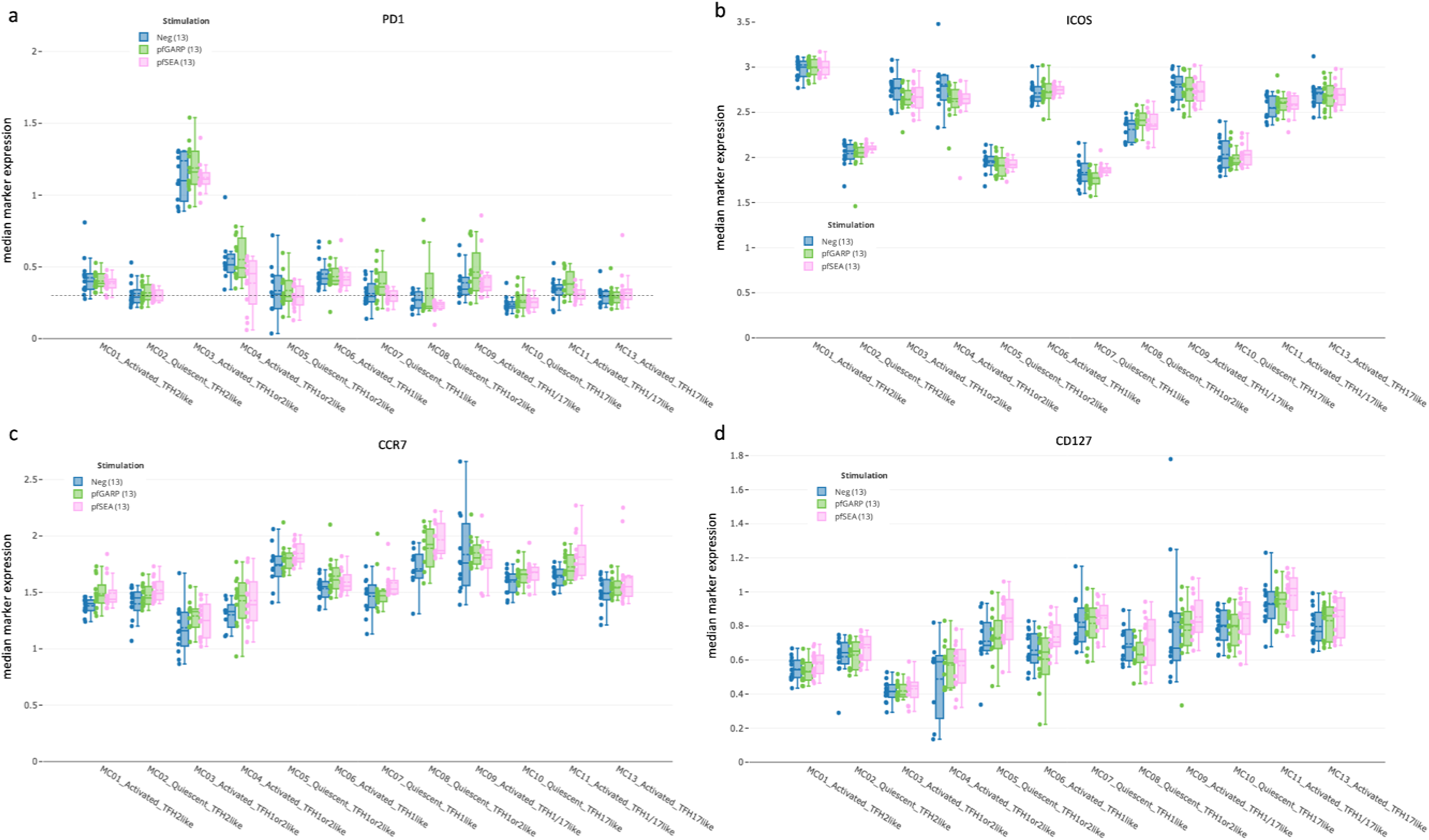
cT_FH_ subset activation state determined by PD1, ICOS, CCR7, and CD127. Box-plots showing **(a)** PD1, **(b)** ICOS, **(c)** CCR7, and **(d)** CD127 expression across all cT_FH_- like meta-clusters from children (n=13) with no stimulation (blue), and after *in vitro* stimulation with *Pf*GARP (green) or *Pf*SEA-1A (pink). Fifty percent of the data points are within the box limits, the solid line indicates the median, the dashed line indicates the mean, and the whiskers indicate the range of the remaining data with outliers being outside that range.

We also observed additional nuances in the expression pattern for other cT_FH_1-like meta-clusters. The cT_FH_1-like MC06 cluster had an activated/effector profile, whereas the cT_FH_1-like MC07 cluster had a quiescent/effector profile. The cT_FH_1or2-like MC05 and MC08 clusters were defined as quiescent/memory (overall high CCR7 expression) yet with very low expression of ICOS for the MC05 cluster.

Likewise, cT_FH_2-like and cT_FH_17-like meta-clusters also displayed heterogeneity. The cT_FH_2-like MC01 cluster had an activated/effector profile, whereas the cT_FH_2-like MC02 cluster seemed to be a quiescent/effector subset. The cT_FH_1/17-like subsets MC09 and MC11 overall had an activated profile, although they had higher expression of CCR7 and CD127 compared to other subsets, suggesting an activated/memory phenotype. Finally, the cT_FH_17-like MC10 cluster seemed quiescent, whereas cT_FH_17-like MC13 had an activated/effector profile. Of note, because of our short stimulation time (6 hrs), we were unable to find statistical differences in the CD40L expression between groups as only few individuals responded through it (Supplemental Figure 7). However, our analysis methods revealed a higher degree of previously unrecognized heterogeneity within circulating cT_FH_ cells.

### Activated cT_FH_1or2-, cT_FH_1-, and quiescent cT_FH_1or2-like subsets were more abundant in children

After having deconvoluted cT_FH_ cells into 12 subsets, we next wanted to determine whether their abundance differed by age or after antigen stimulation. Using Uniform Manifold Approximation and Projection (UMAP) visualization, we found that cT_FH_ dimensional reduction was contiguous as meta-clusters merged with each other; in addition, there were notable differences between adults and children (Figure 5a). We found that activated PD1^high^ cT_FH_1or2-like (MC03), activated cT_FH_1-like (MC06), and quiescent ICOS^high^ cT_FH_1or2-like (MC08) subsets were significantly more abundant in children compared to adults regardless of the stimulation conditions (Figure 5b, 5c, and 5d, *p*<0.05). In contrast, the quiescent *Pf*SEA-1A- and *Pf*GARP-specific cT_FH_2-like cluster (MC02) was significantly more abundant in adults compared to children (Figure 5c and 5d, *p*<0.05). Interestingly, following *Pf*GARP stimulation, the activated cT_FH_1/17-like subset (MC09) became more abundant in children compared to adults (Figure 5d*, p*<0.05 with a False Discovery Rate=0.08), but no additional subsets shifted phenotype after *Pf*SEA-1A stimulation (Figure 5c). Of note, the activated PD1^low^ cT_FH_1or2-like cells (MC04) seemed more abundant in non-stimulated adults and *Pf*GARP-stimulated children, but, because these observations were not present in all the participants, they did not achieve statistical significance in EdgeR.

**Figure 5:**
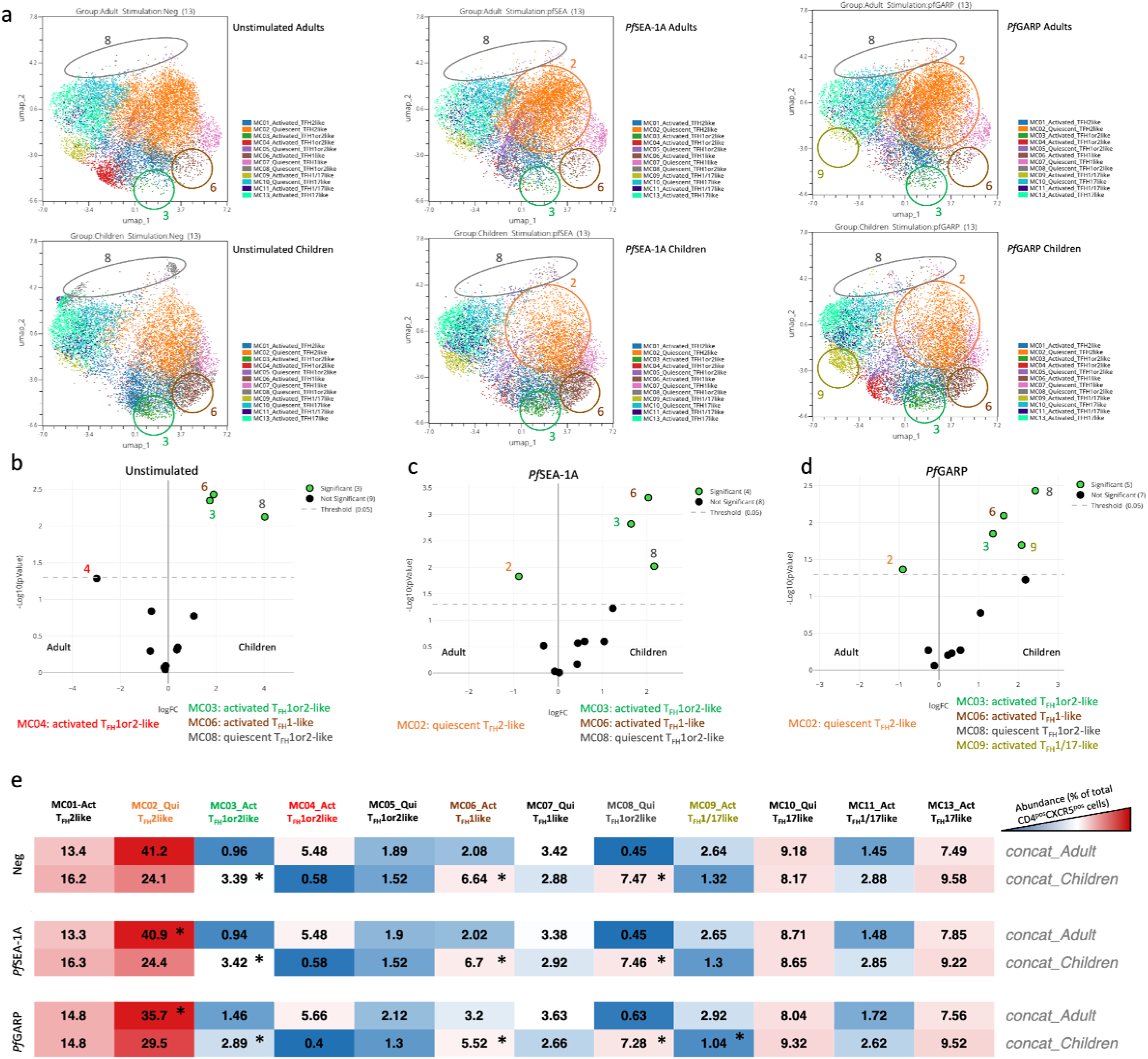
Differences in abundance of antigen-specific cT_FH_ meta-clusters for adults and children. **(a)** UMAP plots showing the 12 different cT_FH_ meta-clusters in adults (top three plots, n=13) and in children (bottom three plots, n=13) in the absence of stimulation or after *in vitro* stimulation with *Pf*SEA-1A or *Pf*GARP, from left to right, respectively. Colored circles highlight the meta-clusters showing differences in their abundance between adults and children for each condition. An EdgeR statistical plot was performed to assess the change in abundance of the 12 meta-clusters between adults and children after **(b)** no stimulation or stimulation with **(c)** *Pf*SEA-1A or **(d)** *Pf*GARP. EdgeR plots indicate which meta-clusters are significantly abundant between two groups by using green color dots. The Y-axis is the - log_10_(*p*-value) and the X-axis is the log(FC). Green dots were statistically significant (*p*<0.05). Numbers next to the dots indicate the meta-cluster. **(e)** An abundance heatmap indicates the percentage (black numbers) of each meta-cluster within the total number of CD3^pos^CD4^pos^CXCR5^pos^CD25^neg^ cells for adults and children (concatenated from 13 participants in each group) under the different conditions: no stimulation (Neg) or stimulation with *Pf*SEA-1A or *Pf*GARP. The color scale ranges from high expression (red) to low/no expression (blue). The star in the heatmap indicates which meta-cluster is significantly abundant in children or adults based on the EdgeR results.

The abundance heatmap (Figure 5e) reiterates the differences observed between children and adults and highlights important considerations when assessing the potential role of each cT_FH_ subset in assisting with cognate antibody production. Overall, the most common cT_FH_ subset in both children and adults is the quiescent cT_FH_2-like cells (MC02, 24.1% and 41.2% respectively). However, the antigen-specific differences in the cT_FH_ subset abundance for children (MC09 for *Pf*GARP) and for adults (MC02 for both *Pf*SEA-1A and *Pf*GARP) suggest that children engage different cT_FH_ cells as they are developing immunity. Of note, only the activated PD1^low^ cT_FH_1or2-like cells (MC04) was less abundant in children with low compared to high HRP2 antibody levels (Supplemental Figure 8), suggesting that this subset may be involved in short-term antibody production. Overall, this comprehensive examination of the abundance of cT_FH_ subsets demonstrates important diversity based on age and malaria-antigen specificity.

### PfSEA-1A and PfGARP induced IL-4, Bcl6, and cMAF from a broad range of cT_FH_ subsets in children

Because the children in this cohort were 7 years of age and resided in a malaria-holoendemic area, they had ample time to develop premunition. To assess antigen-specific cytokine and transcription factor expression signatures and further characterize cT_FH_ subsets, we generated clustered heatmaps of the median fluorescence intensity (MFI) of each analyte (IFN, IL-4, IL-21, Bcl6, and cMAF) for children (Supplemental Figure 9a) and adults (Supplemental Figure 9b). Next, using these MFI data, we performed Wilcoxon paired two-tailed t-tests to compare *Pf*SEA-1A and *Pf*GARP stimulation to unstimulated cells (Figure 6a and 6b, respectively). Significant differences in these expression profiles allowed us to further characterize meta-clusters into three main groups. Group 1: cT_FH_2-like (activated MC01 and quiescent MC02) and activated cT_FH_1or2-like (PD1^high^MC03 and PD1^low^MC04); Group 2: activated and quiescent cT_FH_1-like (MC06 and MC07); Group 3: activated cT_FH_1/17-like (MC11) and cT_FH_17-like (quiescent MC10 and activated MC13). For children, *Pf*SEA-1A and *Pf*GARP induced robust IL-4 expression in 9 out of 12 cT_FH_ meta-clusters (*p*-values ≤ 0.0105); although the composition of which cT_FH_ subsets were engaged differed slightly by antigen (quiescent cT_FH_1or2-like ICOS^low^ MC05 versus quiescent cT_FH_2-like MC02, respectively). In contrast to IL-4, we observed no change in expression for IFN□ or IL-21 after *in vitro* antigen stimulation (except for quiescent cT_FH_1or2-like ICOS^high^ MC08, *p*- value=0.0391), suggesting that these cytokines are not informative to define the development of antigen-specific cT_FH_ subsets in children.

**Figure 6:**
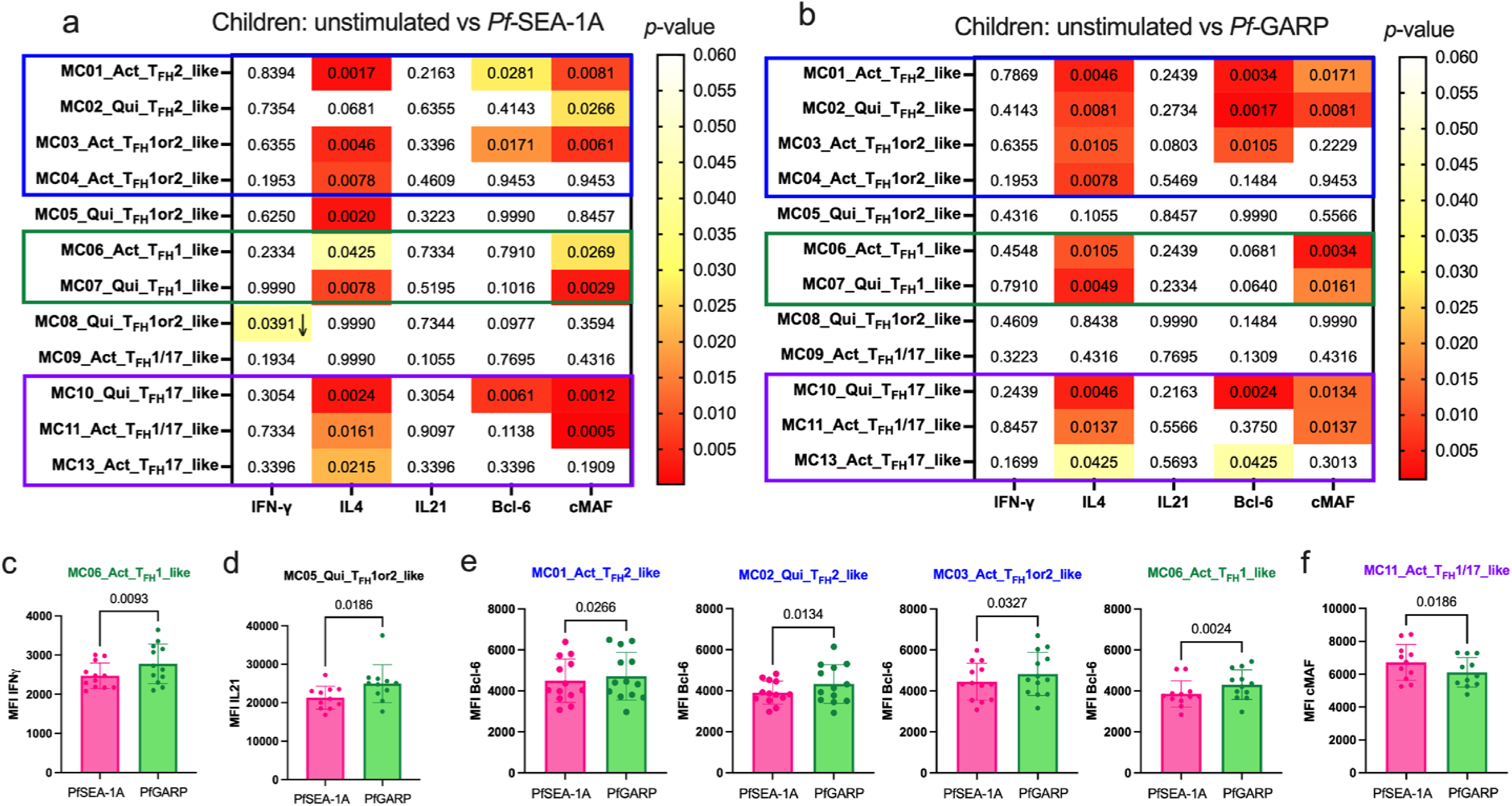
Diverse *Pf*-malaria antigen-specific expression patterns of cT_FH_-defining cytokines and transcription factors for children. Heatmaps of Wilcoxon-paired two-tailed t-test *p*-values are shown for cT_FH_ meta-clusters comparing **(a)** *Pf*SEA-1A and **(b)** *Pf*-GARP versus unstimulated PBMCs from children (n=13) for each cytokine (IFNγ, IL-4, and IL-21) and transcription factor (Bcl6 and cMAF). The color scale indicates the significance of the *p*- value: white (non-significant, *p*>0.05), yellow (0.05>*p*>0.02), orange (0.02>*p*>0.005), and red (highly significant, *p*<0.005). The down arrow indicates a decrease of expression from unstimulated to stimulated condition, whereas no arrow indicates an increase of expression from unstimulated to stimulated condition. The cT_FH_ meta-clusters that co-expressed transcription factors were grouped as follows: Group 1 (blue line), Group 2 (green line), and Group 3 (purple line). Bar plots indicating mean with SD of the median intensity fluorescence (MFI) of **(c)** IFNγ, **(d)** Bcl6, and **(e)** cMAF for the cT_FH_ meta-clusters showing significant statistical differences between *Pf*SEA-1A and *Pf*GARP stimulations. The *p*-values from Wilcoxon-paired two-tailed t-tests are indicated.

We found that *Pf*SEA-1A (Figure 6a) and *Pf*GARP (Figure 6b) induced similar Bcl6 and cMAF expression profiles from some of the same cT_FH_ subsets: both transcription factors were expressed by activated cT_FH_2-like MC01 and quiescent cT_FH_17-like MC10, but only cMAF was expressed in activated and quiescent cT_FH_1-like MC06, MC07, and activated cT_FH_1/17-like MC11 subsets. However, *Pf*SEA-1A induced Bcl6 and cMAF from activated PD1^high^MC03, whereas *Pf*GARP only induced Bcl6. In contrast, the quiescent cT_FH_2-like MC02 subset did not seem to respond to *Pf*SEA-1A (only cMAF was significant, *p*=0.0266, Figure 6a, Supplemental Figures 10 and 11), whereas *Pf*GARP stimulation induced significantly more IL-4, Bcl6, and cMAF compared to unstimulated cells (*p*=0.0081, *p*=0.0017, and *p*=0.0081, respectively, Figure 6b, Supplemental Figures 10 and 11). We then compared the response intensity between *Pf*SEA-1A and *Pf*GARP stimulations and found significant differences in IFN□, IL-21, Bcl6, and cMAF expression levels (Figure 6c, 6d, 6e, and 6f, respectively). In Group 1, Bcl6 expression was significantly higher within activated and quiescent cT_FH_2-like subsets (MC01 and MC02) as well as activated PD1^high^ cT_FH_1or2-like (MC03) cells after *Pf*GARP compared to *Pf*SEA-1A stimulation (*p*=0.0266, *p*=0.0134 and *p*=0.0327, respectively, Figure 6e). Within Group 2, IFN□ and Bcl6 were highly expressed by activated cT_FH_1-like (MC06) after *Pf*GARP stimulation compared to *Pf*SEA-1A stimulation (*p*=0.0093 and *p*=0.0024, respectively, Figure 6c and 6e). Finally, within Group 3, the activated cT_FH_1/17-like cells (MC11) expressed higher cMAF after *Pf*SEA-1A stimulation compared to *Pf*GARP (p=0.0186, Figure 6f). Interestingly, *Pf*GARP induced significantly more IL-21 within the quiescent ICOS^low^ cT_FH_1or2-like (MC05) meta-cluster compared to *Pf*SEA-1A stimulation (*p*=0.0186, Figure 6d). Non-significant differences in cytokines and transcription factors expressed by cT_FH_ subsets between conditions are shown in Supplemental Figures 12 and 13, respectively.

### PfGARP induced IL-4, Bcl6, and cMAF expression in activated cT_FH_1/17- and cT_FH_17-like subsets in adults

A similar heatmap was generated for adult expression profiles comparing *Pf*SEA-1A and *Pf*GARP stimulated to unstimulated cells (Figure 7). Here, we found that both *Pf*SEA-1A and *Pf*GARP induced significant expression of both IFN□ and IL-4 for activated (MC01) and quiescent (MC02) cT_FH_2-like cells (Figure 7a and 7b, Supplemental Figure 12). Whereas *Pf*GARP also induced IL-4 and IFN□ expression from quiescent and activated cT_FH_17-like cells (MC10 and MC13), in addition to Bcl6 and cMAF for MC13 (Figure 7b, Supplemental Figure 13). This observation was surprising because IFN□ expression is commonly used to categorize the cT_FH_1 subset (Group 2); however, as shown earlier, quiescent and activated cT_FH_17-like cells (MC10 and MC13) did not express CXCR3 (Figure 3d and 3e). In contrast, activated the cT_FH_1/17-like cells (MC11) only responded to *Pf*GARP (Figure 7b, Supplemental Figure 12), expressing higher levels of IL-4, IL-21, Bcl6, and cMAF. Finally, while assessing the differences between the two malaria antigens, we found that *Pf*GARP induced more Bcl6 expression than *Pf*SEA-1A within the quiescent cT_FH_2-like subset (MC02, *p*=0.0327) and the quiescent ICOS^low^ cT_FH_1or2-like cells (MC05, *p*=0.0105), as well as within the quiescent cT_FH_17-like subset (MC10, *p*=0.0479) and the activated cT_FH_1/17-like cells (MC11, *p*=0.0020) (Figure 7d). The MC11 cells also expressed higher IL-21 levels after *Pf*GARP compared to *Pf*SEA-1A stimulation (*p*=0.0273, Figure 7c); whereas *Pf*SEA-1A induced cMAF within the activated cT_FH_2-like subset (MC01, *p*=0.0134) but *Pf*GARP did not (Figure 7e). Overall, the main observation for adults is that *Pf*SEA-1A predominantly induced IL-4 from slightly more than half of the cT_FH_ clusters, whereas *Pf*GARP induced a broader range of cytokines and transcription factors but within the activated cT_FH_1/17-like cells (MC11) and quiescent and activated cT_FH_17-like subset (MC10 and MC13) similar to children. This analysis shows clear differences in cT_FH_ subset specificity by malaria antigen and cT_FH_ subset engagement by age group, with the cT_FH_ repertoire becoming more restricted in adults compared to children.

**Figure 7:**
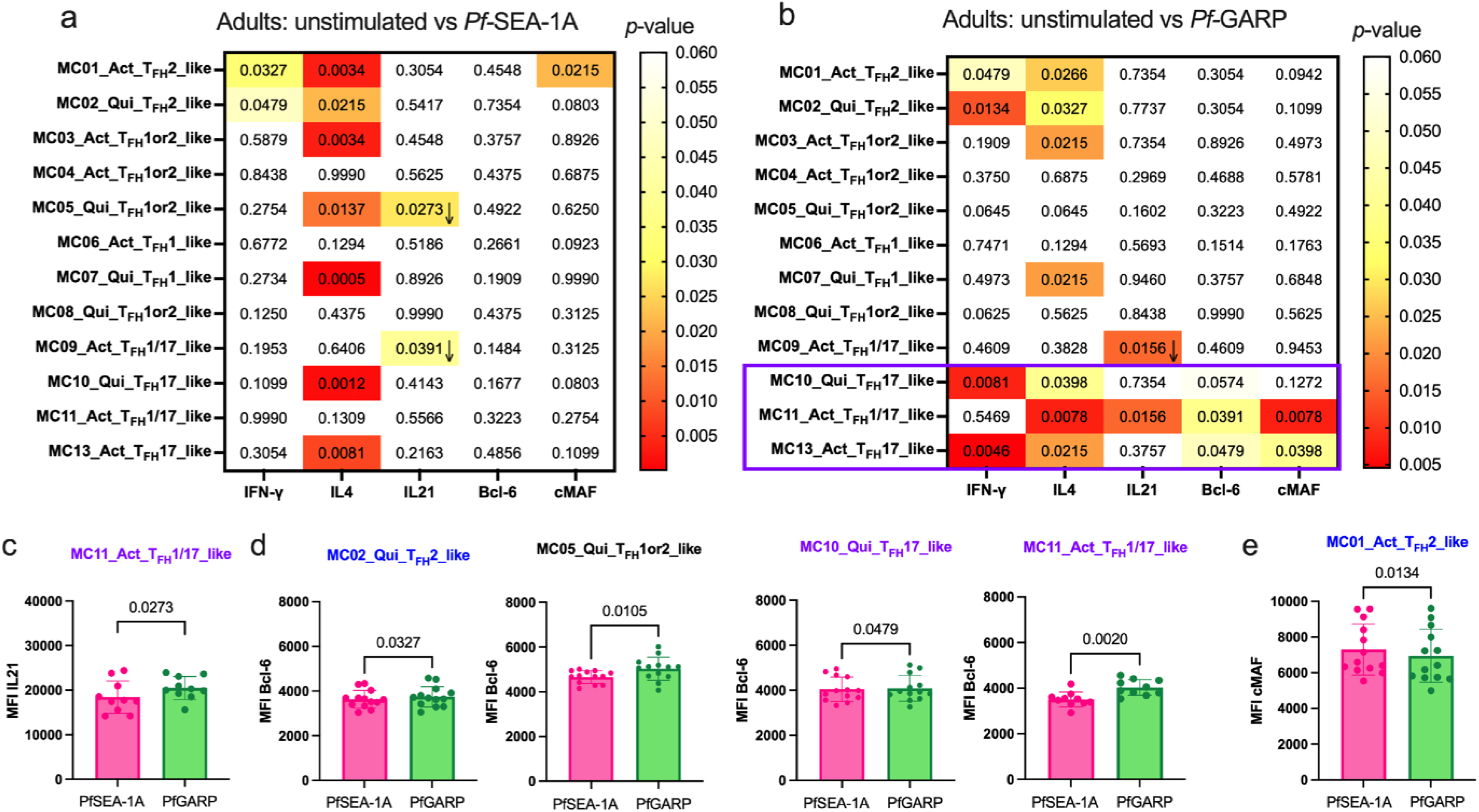
Limited *Pf*-malaria antigen-specific expression patterns of cT_FH_ defining cytokines and transcription factors for adults. Heatmaps of Wilcoxon-paired two-tailed t- test *p*-values are shown for cT_FH_ meta-clusters comparing **(a)** *Pf*SEA-1A or **(b)** *Pf*-GARP versus unstimulated PBMCs from adults (n=13) for each cytokine (IFNγ, IL-4, and IL-21) and transcription factors (Bcl6 and cMAF). The color scale indicates the significance of the *p*-value: white (non-significant, *p*>0.05), yellow (0.05>*p*>0.02), orange (0.02>*p*>0.005), and red (highly significant, *p*<0.005). The down arrow indicates a decrease of expression from unstimulated to stimulated condition, whereas no arrow indicates an increase of expression from unstimulated to stimulated condition. The only cT_FH_ meta-clusters that expressed transcription factors were in Group 3 (purple box). Bar plots indicate the mean with SD of the median fluorescence intensity (MFI) of **(c)** Bcl6 and (**d**) cMAF for the cT_FH_ meta-clusters showing significant statistical differences between *Pf*SEA-1A and *Pf*GARP stimulations. The *p*-values from Wilcoxon-paired two-tailed t-tests are indicated.

### The activated cT_FH_1or2-like subset is more abundant in participants with high anti-PfGARP antibodies

As shown in Figure 1b, a broad range of anti-*Pf*GARP IgG antibody levels were found in both children and adults. Thus, we wanted to determine whether the abundance of any of the cT_FH_ subsets was associated with the level of anti-*Pf*GARP IgG antibodies. When stratifying by high versus low anti-*Pf*GARP IgG antibody levels, we found that an activated cT_FH_1or2-like subset (MC04) was more abundant in participants with high levels of anti-*Pf*GARP IgG for both children (Figure 8a) and adults (Figure 8b) after *Pf*GARP stimulation compared to participants with low levels or an absence of anti-*Pf*GARP IgG. However, even though the *p*- values were significant for both children and adults (*p*=0.02 and *p*=0.004, respectively), the FDR was less than 0.05 only for the adults (FDR=0.018). This suggests that this particular subset might be important for the generation of anti-*Pf*GARP antibodies. Interestingly, activated cT_FH_1-like (MC06) and quiescent cT_FH_1-like (MC07) cells were more abundant for adults with low or no anti-*Pf*GARP IgG antibodies compared to those with high levels (*p*=0.0024 with FDR=0.0185 and *p*=0.0034 with FDR=0.0185, respectively) consistent with previous observations describing T_FH_1 subsets as inefficient help for antibody production^29^.

**Figure 8:**
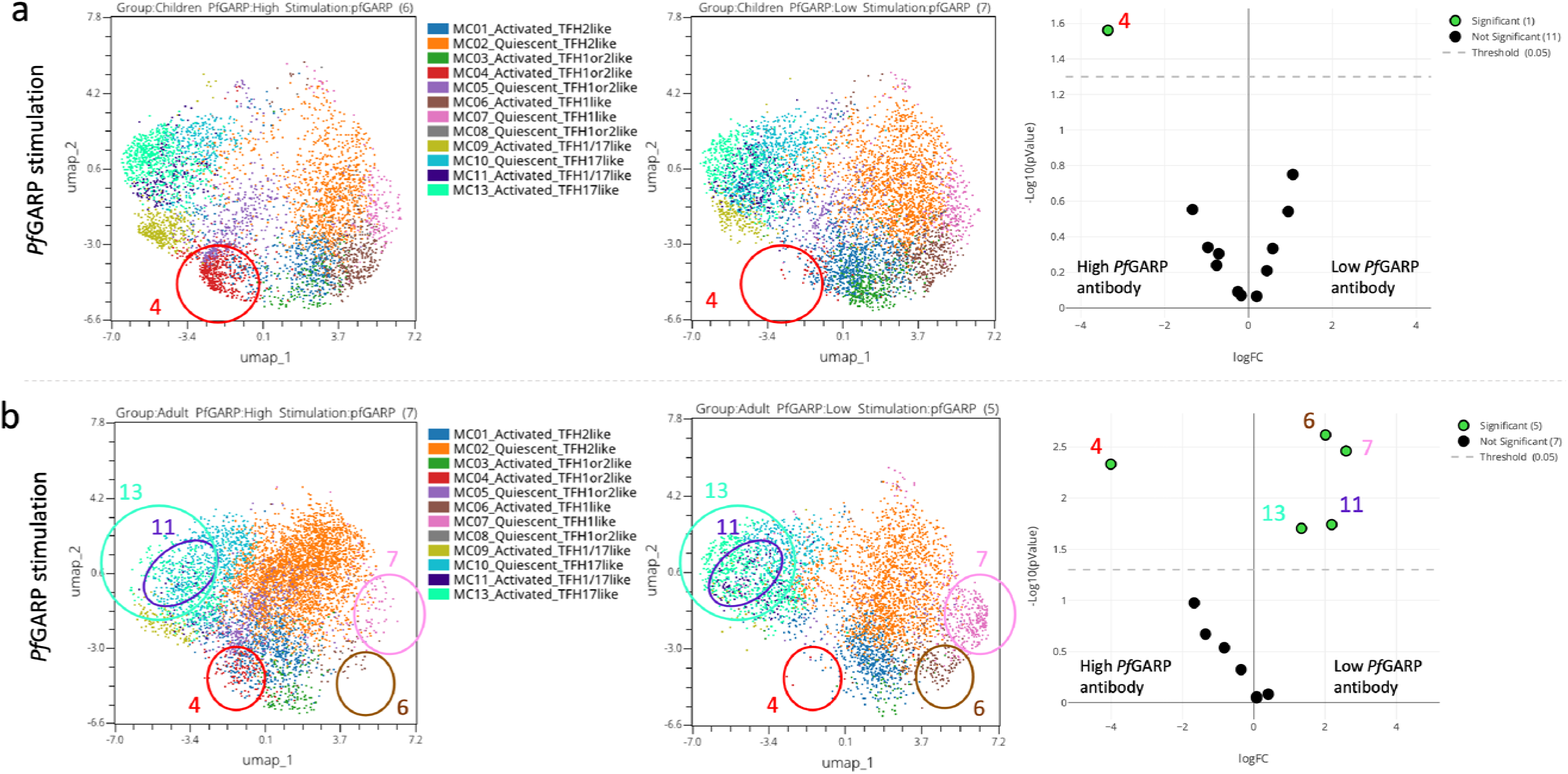
Abundance of cT_FH_ meta-clusters stratified by anti-*Pf*GARP IgG antibody levels. UMAP and EdgeR analyses of **(a)** children (n=13) and **(b)** adults (n=13) where significant differences (*p*<0.05) in the abundance of the cT_FH_ meta-clusters are circled on high versus low *Pf*GARP antibody level (left to right) in the UMAP plot and green dots on the volcano plot (far right).

## Discussion

The overall aim of this study was to define cT_FH_ subsets using unbiased clustering analysis and assess their malaria antigen-specific (*Pf*SEA-1A and *Pf*GARP)^6,7,39^ profiles for adults and children residing in a malaria-holoendemic area of Kenya. Contrary to previous publications,^29,30^ our study found that children not only respond via their cT_FH_1-like subsets but also engage a broader spectrum of cT_FH_ subsets. In fact, cytokine and transcription factor profiles for children involved cT_FH_1-, cT_FH_2-, cT_FH_17-, and cT_FH_1/17-like subsets (summarized in Figure 9), whereas in adults, the dominant antigen-specific memory response was from cT_FH_17- and cT_FH_1/17-like subsets but only for *Pf*GARP. Thus, revealing a potential difference in engaging cT_FH_ help between the two malaria vaccine candidates evaluated in this study.

**Figure 9:**
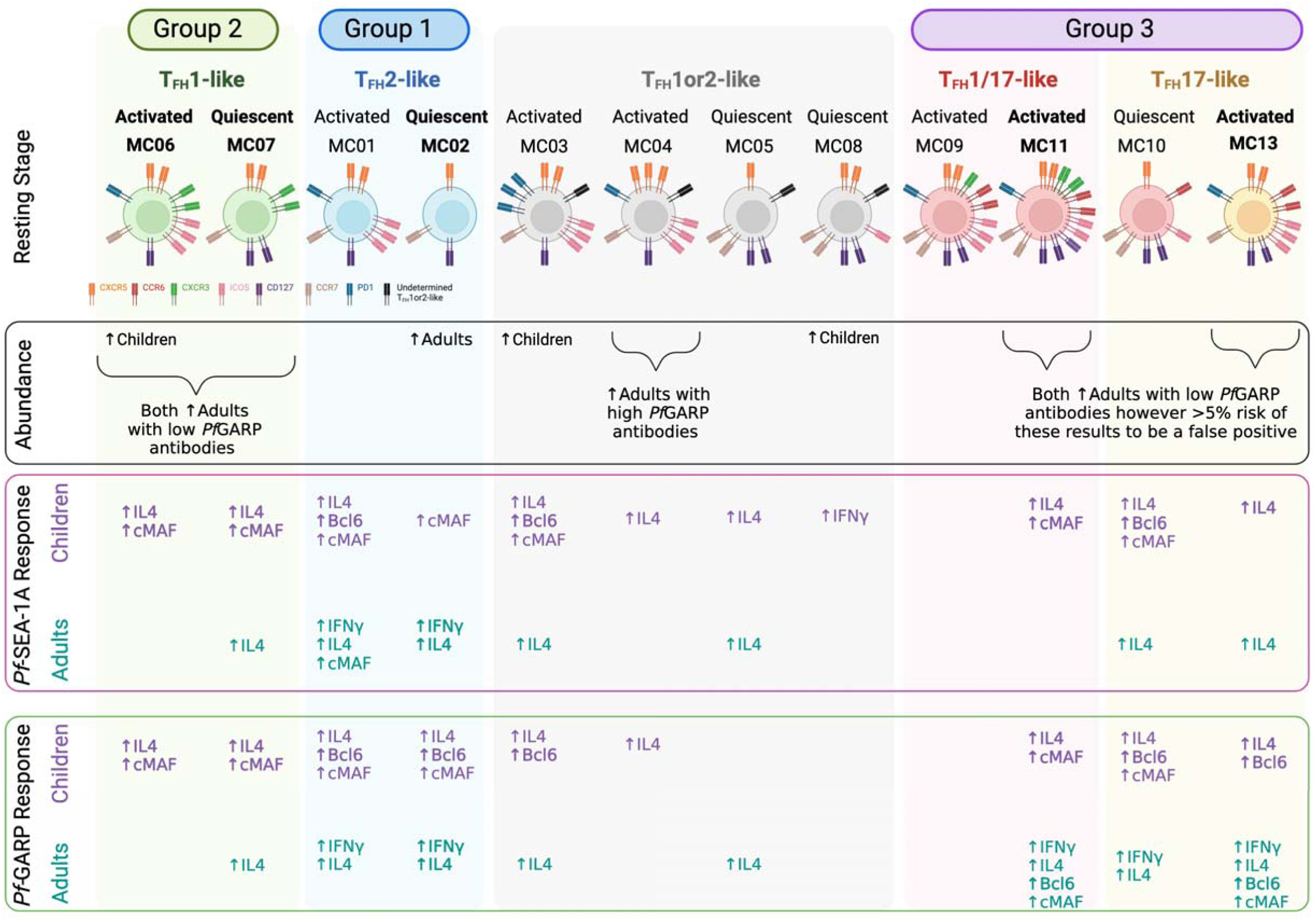
Summary. Illustration of combined findings showing remarkable differences in cT_FH_ subset abundance and *Pf*-malaria antigen-specific cytokine and transcription factor responses between children and adults residing in a malaria-holoendemic region of Kenya. Figure created with BioRender.

Several key points can be made from our results. First, it may be too soon in the field of cT_FH_ biology to select only one or two types of cT_FH_ subset(s) to measure when evaluating malaria vaccine candidates, especially if we miss essential, albeit transient, correlates of protection for children which may differ in adults. Longitudinal studies with clinical outcomes are needed to determine the impact of the changing dynamics of cT_FH_ subset adaptations after repeated malaria infections that lead to premunition and parasite clearance. A Ugandan study by Chan et al.^31^ demonstrated an age-associated change in cT_FH_ subsets independent of malaria and, thus, supports the importance of accounting for age when evaluating antigen-specific cT_FH_ profiles. Second, our study reveals that adults have a robust cT_FH_17- and cT_FH_1/17-like subset response to *Pf*GARP but not to *Pf*SEA-1A. In addition to demonstrating differences between malaria antigens, this finding supports the premise that cT_FH_17 cells are important for maintaining immunological memory,^40^ which has intriguing implications for evaluating malaria vaccine efficacy. Third, we found a correlation between cT_FH_1or2-like (MC04) and high *Pf*GARP antibodies for both children and adults, indicating that a cT_FH_ subset accompanied by high antibody levels could serve as a potential biomarker of protection. More studies are needed to explore this association; however, we postulate that a coalition of cT_FH_ subsets might be engaged to develop long-lived antibody responses and, therefore, categorizing subsets as efficient or inefficient might be context dependent.^29^

As the field of computational immunology and unbiased clustering analyses evolves, it presents an ongoing challenge to meaningfully define cT_FH_ subsets. Here, we intentionally chose to use cT_FH_1or2-like nomenclature for meta-clusters MC03, MC04, MC05, and MC08 (Figure 9) because of their low (but not null) CXCR3 and robust IL-4 expression along with Bcl6 and cMAF). CXCR3 expression leans toward a cT_FH_1-like polarization, whereas IL-4 expression indicates a cT_FH_2-like profile when we follow the most commonly used cT_FH_ classification.^17,21,22,33^ This difference is crucial as cT_FH_2 cells are described as good promoters for functional GC antibodies^28,30^, whereas cT_FH_1 cells are not.^29,30,41^ Therefore, their definitive classification will require a more in-depth investigation. Although we used malaria antigen-specific stimulation, and not a *Pf*-lysate or infected red blood cells (iRBCs) as previously described,^29–31^ adults had significantly more of the quiescent cT_FH_2-like subset (MC02) compared to children who, instead, had significantly more of the activated cT_FH_1- like subset (MC06), the latest being consistent with previous publications,^29,30^ and more PD1^high^ activated and ICOS^high^ quiescent cT_FH_1or2-like subsets (MC03 and MC08, respectively). However, even when classifying the cT_FH_ meta-clusters as cT_FH_1, cT_FH_2, cT_FH_17, or cT_FH_1/17, the most abundant cT_FH_ subset was cT_FH_2 cells for adults (more than 50%) contradicting a previous study showing only ∼20% of cT_FH_2 cells for the same age range.^31^ As suggested by Gowthaman’s T_FH_ model,^27^ T_FH_1 cells are involved in the development of neutralization antibody responses to viruses and bacteria, whereas parasites, such as helminths, lead to a T_FH_2 response. These observations reinforce the need to assess T_FH_ subset profiles stratified by exposure to potentially immune-modulating co-infections.

Transcription factors cMAF and Bcl6 play essential roles in T_FH_ development and function.^24,42,43^ cMAF induces the expression of various molecules, such as ICOS, PD1, CXCR5, IL-4, and IL-21, which are all essential for T_FH_ function.^42^ It was, therefore, not surprising to find that cMAF significantly increased after antigen stimulation for most of the cT_FH_ subsets in children. However, again, this expression profile was only observed in the activated cT_FH_1/17-like subset (MC11) after *Pf*GARP stimulation in cells from adults, supporting a role for T_FH_17 cells in the maintenance of immunological memory.^40^ Importantly, cMAF cooperates with Bcl6 in T_FH_ development and function and is essential to establish an efficient GC response.^24,44^ Bcl6 is also described as being expressed by mature T_FH_ cells, inducing the expression of CXCR5 and PD1, but it can also regulate IFNγ and IL-17 production.^12,24,44,45^ In our study, *Pf*GARP induced a highly significant increase of Bcl6 in cT_FH_2-like and cT_FH_17-like subsets in children, suggesting that *Pf*GARP may be a better candidate to trigger an efficient humoral response in children compared to *Pf*SEA-1A. Of note, Blimp1, a Bcl6 antagonist,^45^ was absent from our flow panel and would be of interest to assess in future studies.

There are several limitations of human immune profiling studies. Similar to other such studies, we used peripheral blood and, thus, were only able to provide a snapshot of the cT_FH_ cells. Study participants did not have blood-stage malaria infections at the time of blood collection, and children were 7 years of age; thus, our profiles were by design meant to reflect cT_FH_ memory recall responses and demonstrate differences between children and adults. As this was not a birth cohort study design, the number of cumulative malaria infections could have confounded the association between malaria and cT_FH_ subsets, causing a certain cT_FH_ meta-cluster to arise. Most human immunology studies are unable to assess tissue-resident T_FH_ or T_FH_ in the lymph nodes. Therefore, we can only speculate on the composition of T_FH_ cells observed in children that were perhaps short-lived helper cells and not maintained as T cell memory cells or may have trafficked from the blood into tissues over time.^46^ Using an *in vivo* non-human primate model, Potter et al., showed an entry rate of lymphocyte subsets into peripheral lymph nodes per hour of 1.54% and 2.17% for CD4^pos^ central memory (CD45^neg^CCR7^pos^) and effector memory (CD45^neg^CCR7^neg^) T cells, respectively,^46^ both of which may include central and effector memory cT_FH_ cells. With this in mind, cT_FH_ cells seem to be at a crossroads with multiple possible fates. The first one being the recirculation of cT_FH_ cells from lymph node to lymph node, whereby a peripheral blood sampling captures migratory cT_FH_ subsets, which could explain the cT_FH_ subsets we observed still expressing cMAF and Bcl6. A decade ago, it was hypothesized that once a T_FH_ cell leaves the lymph node, it can become a PD1^low^ memory cT_FH_ cell with the possibility of returning to a GC T_FH_ stage after a secondary recall response.^12^ Other possible fates include becoming a PD1^neg^ memory cT_FH_ with the same path after recall or progressing to a non-T_FH_ cell.^12^ More recently, a few studies showed circulating CXCR5^neg^CD4^pos^ cells with T_FH_ functions, such as production of IL-21 and the capability to help B cells in systemic lupus erythematosus and HIV-infected individuals,^47,48^ and importantly researchers mapped these circulating CXCR5^neg^CD4^pos^ cells to an original lymph node CXCR5^pos^ T_FH_ subset,^48^ suggesting another possible outcome for T_FH_ cells outside secondary lymphoid organs. To date, no clear destiny has been established for cT_FH_ cells in humans, thus, highlighting the importance of including a complete panel of cT_FH_ subsets to continue to improve our understanding of their respective roles against different pathogens and eliciting long-lived vaccine-induced antibody responses.

Other limitations of this study include not measuring other cytokines (i.e., IL-5, IL-13, and IL-17) and transcription factors (i.e., T-bet, BATF, GATA3, and RORγt) that have been used in other studies to fully characterize cT_FH_ subsets^12,27^. Although the heterogeneity in the response of CD40L and IFNγ suggests that our tested malaria antigens did not induce significant differences in the expression of these markers in all our participants, our panel did not include other activated induced markers, such as OX40, 4-1BB, and CD69. However, even with our small sample size, we demonstrated significant age-associated differences in malaria antigen-specific responses from different cT_FH_ subsets. To minimize false-positive results that can arise when using algorithms for computational analyses, we ran the statistical tests in triplicate and the clustering algorithm in duplicate, to validate our findings.

In summary, our study provides additional justification for the resources needed to conduct cellular immunological studies of cT_FH_ cell signatures and provides insight into which types of cT_FH_ subsets during a person’s lifespan assist B cells in the GC to produce long-lived plasmocytes and functional antibodies^49^ against malaria. This is particularly important when selecting immune correlates of protection that could be used to predict the efficacy of the next generation of *Pf*-malaria vaccine candidates within various study populations.

## Methods

### Study populations and ethical approvals

Adults and children were recruited from Kisumu County, Kenya, which is holoendemic for *Pf*-malaria. Written informed consent was obtained from each adult participant and every child’s guardian. An abbreviated medical history, physical examination, and blood film were used to ascertain health and malaria infection status at the time of blood sample collection. Participants were also life-long residents of the study area with an assumption that they naturally acquired immunity to malaria. This study was conducted before the implementation of any malaria vaccines. Participants were eligible if they were healthy and not experiencing any symptoms of malaria at the time venous blood was collected. For this cross-sectional immunology study, we selected fourteen 7-year-old children from a larger age-structured prospective cohort study (enrollment age range 3–7 years) and fifteen Kenyan adults. We selected 7-year-olds because of the age-dependent shift in major cT_FH_ subsets occurring after 6 years of age^31^, and as a comparable age published by other studies^29^, to maximize our ability to measure antigen-specific differences in T_FH_ subsets.

Ethical approvals were obtained from the Scientific and Ethics Review Unit (SERU) at the Kenya Medical Research Institute (KEMRI) reference number 3542, and the Institutional Review Board at the University of Massachusetts Chan Medical School, Worcester, MA, USA, IRB number H00014522. Brown University, Providence, RI, USA signed a reliance agreement with KEMRI.

### Plasma and peripheral blood mononuclear cell (PBMC) isolation

Venous blood was collected in sodium heparin BD Vacutainers and processed within 2 hours at the Center for Global Health Research, KEMRI, Kisumu. Absolute lymphocyte counts (ALC) were determined from whole blood using BC-3000 Plus Auto Hematology Analyzer, 19 parameters (Shenzhen Mindray Bio-Medical Electronics Co.). After 10 mins at 1,000*g* spin, plasma was removed and stored at −20 °C, and an equivalent volume of 1X PBS was added to the cell pellet. PBMCs were then isolated using Ficoll-Hypaque density gradient centrifugation on SepMate (StemCell). PBMCs were frozen at 5×10^6^ cells/ml in a freezing medium (90% heat-inactivated and filter-sterilized fetal bovine serum [FBS] and 10% dimethyl sulfoxide [DMSO, Sigma]) and chilled overnight in Mr. Frosty™ containers at −80 °C before being transferred to liquid nitrogen. For transport to the USA, an MVE vapor shipper (MVE Biological Solutions) was used to maintain the cold chain.

### In-vitro stimulation assay

PBMCs were thawed in 37 °C filtered-complete media (10% FBS, 2 mM L-Glutamine, 10 mM HEPES, 1X Penicillin/Streptomycin) and spun twice before resting overnight in a 37 °C, 5% CO_2_ incubator. PBMCs were counted using Trypan Blue (0.4%) and a hemocytometer, and the cell survival was calculated. Our samples showed a median of 94.6% live cells (25% percentile of 92%; 75% percentile of 97%). Using a P96 U-bottom plate, 1×10^6^ PBMCs per well were placed in culture with one of the following stimulation conditions: *Pf*SEA-1A^6^ (5 µg/ml) or *Pf*GARP^7^ (10 µg/ml) both produced in the Kurtis lab (Brown University); SEB (1 µg/ml; EMD Millipore) was used as a positive control; sterile water (10 µl, the same volume used to reconstitute *Pf*SEA and *Pf*GARP) was used as a negative control. A pool of anti-CD28/anti-CD49d (BD Fast-Immune Co-Stim following the manufacturer’s instructions), GolgiSTOP (0.7 µg/ml), and GolgiPLUG (0.1 µg/ml) (BD Biosciences) were added to each well before incubating cells at 37 °C for 6 hours.

### Cell staining and flow cytometry

A multiparameter spectral flow cytometry panel was used to characterize cT_FH_ cell subsets: CCR6-BV421 (RRID: AB_2561356), CD14-Pacific Blue (RRID: AB_830689), CD19-Pacific Blue (RRID: AB_2073118), CCR7-BV480 (RRID: AB_2739502), IFNγ-BV510 (RRID: AB_2563883), CD127-BV570 (RRID: AB_10900064), CD45RA-BV605 (RRID: AB_2563814), PD1-BV650 (RRID: AB_2738746), CXCR3-BV711 (RRID: AB_2563533), CD25-BV750 (RRID: AB_2871896), CXCR5-BV785 (RRID: AB_2629528), Bcl6-AF488 (RRID: AB_10716202), CD3-Spark Blue 550 (RRID: AB_2819985), CD8-PerCP-Cy5.5 (RRID: AB_2044010), IL-21-PE (RRID: AB_2249025), IL-4-PE-Dazzle (RRID: AB_2564036), CD4-PE-Cy5 (RRID: AB_314078), ICOS-PE-Cy7 (RRID: AB_10643411), cMAF-eFluor 660 (RRID: AB_2574388), CD40L-AF700 (RRID: AB_2750053), and Zombie NIR (BioLegend cat# 423106) for Live/Dead staining. Cells were fixed and permeabilized for 45 mins using the transcription factor buffer set (BD Pharmingen) followed by a wash with the perm-wash buffer. Intracellular staining was performed at 4 °C for 45 more mins followed by two washes using the kit’s perm-wash buffer. Data was acquired on a Cytek Aurora with 4-lasers (UMass Chan Flow Core Facility) using SpectroFlo® software (Cytek) and compensation for unmixing and fluorescence-minus-one controls. Quality control of the data was performed using SpectroFlo®, and the multi-parameter analysis was performed with OMIQ data analysis software (www.omiq.ai). Thus, we assessed the expression of markers commonly used to define the following different cT_FH_ (CD4^pos^CD25^neg^CXCR5^pos^) subsets: cT_FH_1-like (CCR6^neg^CXCR3^pos^), cT_FH_2-like (CCR6^neg^CXCR3^neg^), and cT_FH_17-like (CCR6^pos^CXCR3^neg^),^17^ as well as quiescent/central memory cT_FH_ (CCR7^high^PD-1^neg^ICOS^neg^) or activated/effector memory cT_FH_ cells (CCR7^low^PD-1^pos^ICOS^pos^).^17,18^ Representative cytoplots can be found in Supplemental Figure 1.

### Multiplex suspension bead-based serology assay

To measure plasma IgG antibody levels to *Pf*SEA-1A^6^ and *Pf*GARP^7^, we used a Luminex bead-based suspension assay as previously published.^50,51^ In addition, previous *Pf* exposure was determined using recombinant proteins to blood-stage malaria antigens: AMA1, MSP1, HRP2, CelTos, and CSP (gifts from Sheetji Dutta, Evelina Angov, and Elke Bergmann from the Walter Reed Army Institute of Research). Briefly, 100 μg of each antigen or BSA (Sigma), as a background control, were coupled to ∼12×10^6^ non-magnetic microspheres (Bio-Rad carboxylated beads) and then incubated with study participant plasma (spun down 10,000*g* for 10 mins and diluted at 1:100 in the assay dilution buffer) for 2 hrs, followed by incubation with biotinylated anti-human IgG (BD #555785) diluted 1:1000 for 1 hr and streptavidin (BD #554061) diluted 1:1000 for 1 hr following the manufacturer’s instructions. The mean fluorescence intensity (MFI) of each conjugated bead (minimum of 50 beads per antigen) was quantified on a FlexMap3D Luminex multianalyte analyzer (Xponent software). Results are reported as antigen-specific MFI after subtracting the BSA value for each individual since background levels can vary between individuals.

### OMIQ analysis

The fcs files were uploaded into the OMIQ platform after passing quality control under SpectroFlo® (Cytek) where compensation was re-checked. In OMIQ, we arcsinh-transformed the scale to allow downstream analysis and then gated on singlet live lymphocytes and sub-sampled the data to yield 87,712 live lymphocytes per sample. Using only lineage markers CD3, CD4, CD8, CD14, CD19, CXCR5, and CD25, FlowSOM consensus meta-clustering was run on 100 clusters based on the 87,712 live lymphocytes per sample with a comma-separated k-value of 75 and Euclidean distance metric. Using these 75 meta-clusters, we defined subsets of cells based on lineage markers, such as CD3^pos^CD8^pos^, CD3^neg^CD14^pos^CD19^pos^, and CD3^pos^CD4^pos^, and then distinguished CD4^pos^CXCR5^pos^CD25^neg^ (cT_FH_) and CD4^pos^CXCR5^neg^CD25^pos^ (T regulatory [T_reg_] or T follicular regulatory [T_FR_]) subsets. CD25 was used to exclude T_reg_ and T_FR_ cells which share numerous markers with cT_FH_ cells.^52–54^ EmbedSOM dimensional reduction was used to visualize the different groups of cells and EdgeR analysis was run to assess the significance of their differences. A clustering heatmap was used to visualize cytokine expression and transcription factor profiles for each subset.

Focusing on the CD4^pos^CXCR5^pos^CD25^neg^ T_FH_ cells, we ran another FlowSOM analysis based on the 1,000 CXCR5^pos^ cells per sample (two samples from the adult group and one sample from the children group were excluded from the analysis as they had less than 1,000 CXCR5^pos^ cells), using extracellular markers CXCR5, CXCR3, CCR6, ICOS, CCR7, CD45RA, CD127, CD40L, and PD1 enabled the identification of 15 meta-clusters. If the fcs file had more than 1,000 CXCR5^pos^ cells, the down-sampling was done randomly by the OMIQ platform algorithm to select only 1,000 CXCR5^pos^ cells within this specific fcs file. From there, we performed Uniform Manifold Approximation and Projection (UMAP) dimensional reduction, heatmaps, and EdgeR analyses; the latter allowed statistical analysis of the cT_FH_ abundances. To demonstrate the reproducibility of these results, statistical analysis algorithms were run at least three times downstream of the same clustering algorithm and downstream of repeated clustering algorithms. To assess statistical differences in cytokines and transcription factor expression, we exported the statistics dataset from OMIQ containing MFI values from each marker (IFN□, IL-4, IL-21, Bcl6, and cMAF) per cluster and for each sample and stimulation condition. To assess cytokines and transcription factors without bias, we chose to use the total MFI expression per meta-cluster with the assumption that cells with increased production of the desired analyte trigger an increase in the overall meta-cluster MFI compared to unstimulated cells, and if there is no production of the desired analyte, the overall MFI will not differ. However, the percentage of positive IFN□, IL-4, IL-21, Bcl6, or cMAF using manual gating can be found in Supplemental Figures 14, 15, and 16 along with the overlay of the gated positive cells on the CD4^pos^CXCR5^pos^CD25^neg^ UMAP and the cytoplots of the gated positive cells for each meta-cluster (Supplemental Figures 14, 15, and 16).

### Statistical analysis

For this cross-sectional immunology study, we selected both male and female study participants (sex defined at birth). There were fourteen 7-year-old children and fifteen adults. Using GraphPad Prism software (version 7.0), age, sex, ALC, and serological data were compared between adults and children. Because the number of participants within each group was too low to verify the normality of the underlying distributions (adults n=15 and children n=14), we chose to use non-parametric tests, including the Mann-Whitney U test (for unpaired analysis) and Wilcoxon signed-rank test (for paired analysis). When data passed the normality test (D’Agostino and Pearson test), we used Welch’s parametric test. All tests were two-tailed with a *p-value* < 0.05 for significance. Because of the exploratory nature of the analysis, we did not use any adjustment of the *p*-value for multiple comparisons. The tests used are indicated in the legend of each figure. Results were expressed as the mean with standard deviation and exact *p*-values for dot plots.

To perform a statistical analysis of the cytokine and transcription factor expression from each cT_FH_ subset, the exported data file from OMIQ was integrated into GraphPad Prism, and a non-parametric Wilcoxon paired two-tailed *t-*test analysis was done (n=13 in each group). As this analysis generated numerous bar plots (all included in the Supplemental Figures 12 to 15), to better visualize the cytokines and transcription factor patterns, the *p*- values obtained from each analysis are presented using a non-clustering heatmap.

### Role of the funding source

The funders were not involved in the study design, sample collection, data analysis, interpretation of the data, or the writing of the manuscript and its decision to be submitted for publication.

## Contributors

CN, JMO, JK, and AMM designed the research study; CF and SPT conducted experiments; CF, JK, and AMM analyzed data; CF, SPT, JK, and AMM contributed to experimental design; HWW, BO, JMO, JK, and AMM conducted and supervised the clinical study and the sampling; CF generated figures; HWW and BO verified underlying data; CF and AMM led manuscript preparation with feedback from all the authors. All authors have read and approved the final version of the manuscript.

## Data Availability

Deidentified raw data from this manuscript are available from the ImmPort platform under the accession study number SDY2534.

## Declaration of Competing Interest

Dr. Kurtis was the principal investigator on 1R01AI127699-01A1, which supported this study. He holds several patents related to the use of *Pf*SEA-1 and *Pf*GARP as vaccine candidates for *P. falciparum* and has consulted for and is an equity holder in Ocean Biomedical.

## Acknowledgments

The authors would like to thank the children and their families for participating in this study. We thank the field and lab teams for their work collecting data and processing blood samples. We also thank Dr. Melanie Trombly from UMass Chan for proofreading our manuscript. This manuscript was approved for publication by KEMRI.

This study was supported by NIH R01 AI127699 (Kurtis).

## Notes

### Summary of Updates

The updated version of the manuscript has manual gating shown in the supplemental figures 14, 15 and 16 which confirmed our results found with the clustering analysis. We detailed these figures in the methods. We also added references suggested by the eLife reviewers; the limitation of missing activation induced markers (AIM) was added to the discussion; details on the cell fixation for flow cytometry was added to the methods; and the punctuation was revised throughout the entire manuscript.

