## Supplemental figures 1 to 16 for "T follicular helper cell profiles differ by malaria antigen and for children compared to adults"

**Supplementals**

**Supplemental_Figure_1:**

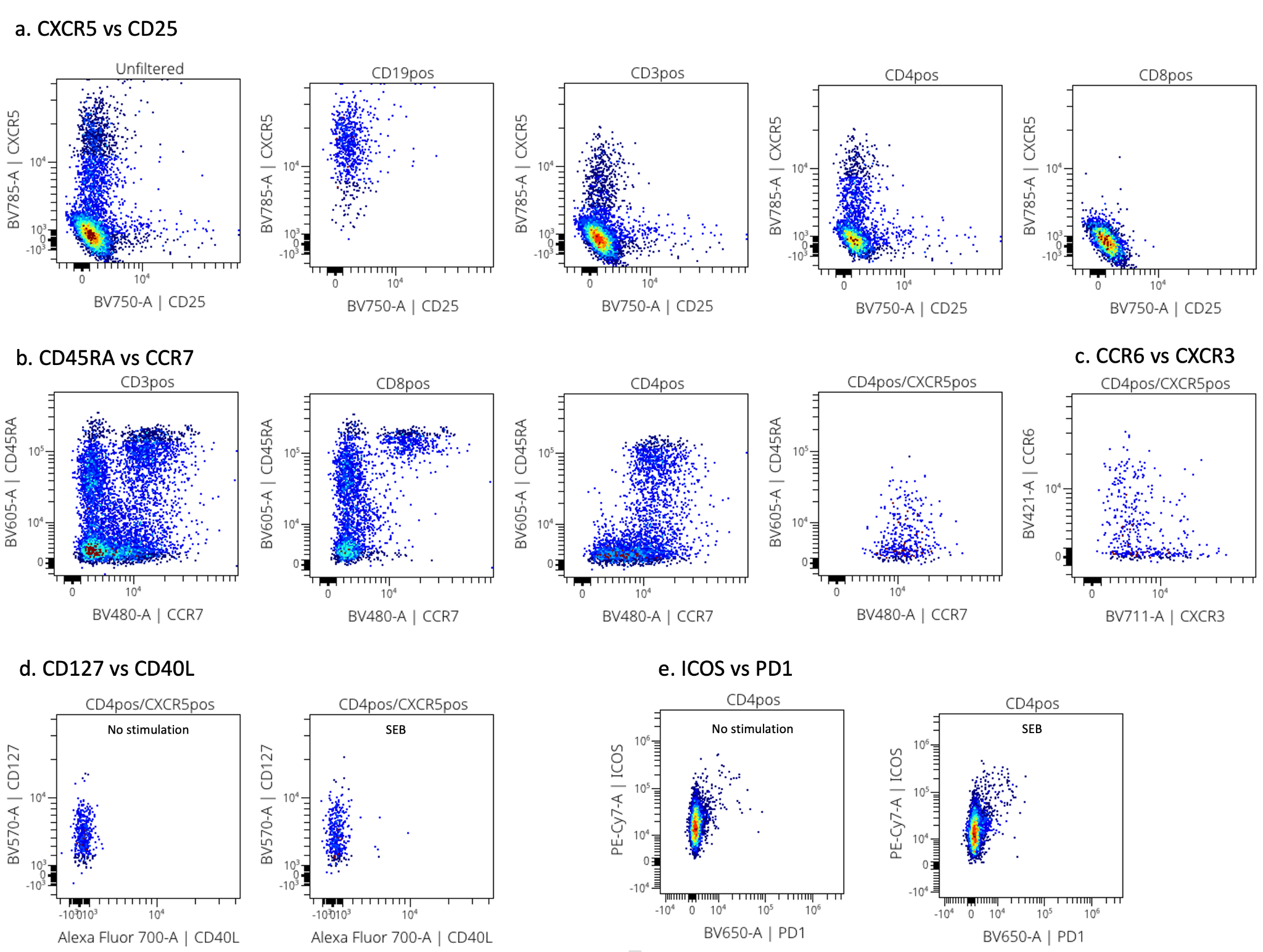

**Supplemental Figure 1: Representative cytoplots of the extracellular flow staining.** Panel **a** shows the CXCR5 vs CD25 cytoplots on unfiltered, CD19^pos^, CD3^pos^, CD4^pos^ and CD8^pos^ cells from left to right. Panel **b** shows CD45RA vs CCR7 cytoplots on CD3^pos^, CD8^pos^, CD4^pos^ and CD4^pos^CXCR5^pos^ cells from left to right. Panel **c** shows CCR6 vs CXCR3 staining on CD4^pos^CXCR5^pos^ cells. Panel **d** shows CD127 vs CD40L expression within CD4^pos^CXCR5^pos^ cells by unstimulated and SEB stimulated cells. Panel **e** shows ICOS vs PD1 cytoplots after no stimulation or SEB 6h stimulation.

**Supplemental_Figure_2:**

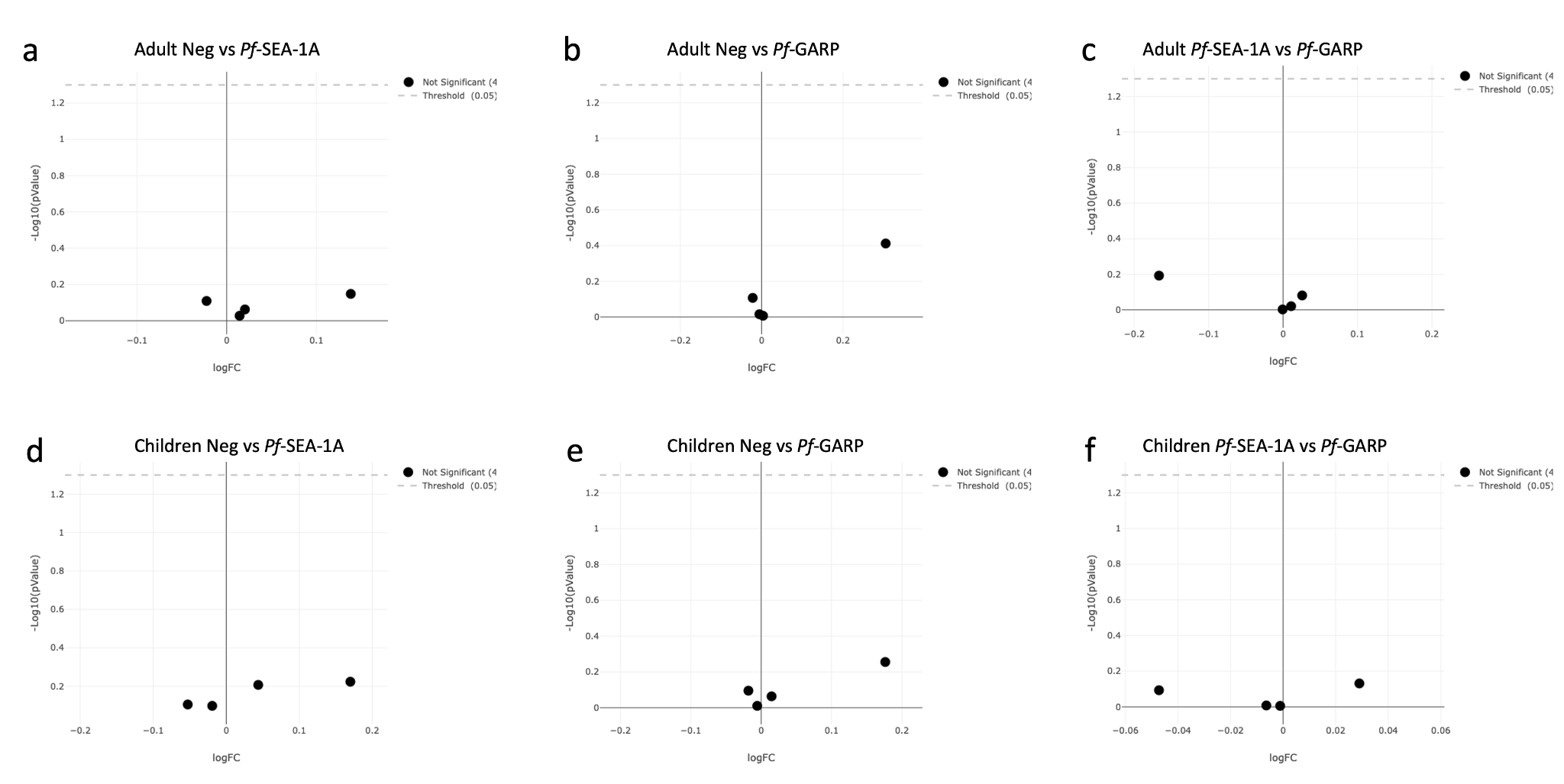

**Supplemental Figure 2: EdgeR analysis of the CD4^pos^, CD4^pos^CXCR5^pos^, CD8^pos^ and CD14/CD19^pos^ populations under different stimulation conditions in adults (n=15) and children (n=14).** Volcano plot from EdgeR analysis comparing CD4^pos^, CD4^pos^CXCR5^pos^, CD8^pos^, and CD14/CD19^pos^ populations from adult PBMCs under the following conditions: **(a)** unstimulated versus *Pf*SEA-1A, **(b)** unstimulated versus *Pf*GARP, and **(c)** *Pf*SEA-1A versus *Pf*GARP. Volcano plot from EdgeR analysis comparing CD4^pos^, CD4^pos^CXCR5^pos^, CD8^pos^,and CD14/CD19^pos^ populations from children PBMCs under the following conditions: **(d)** unstimulated versus *Pf*SEA-1A, **(e)** unstimulated versus *Pf*GARP, and **(f)** *Pf*SEA-1A versus *Pf*GARP. The green dots are statistically significant whereas the black dots are not, here no green dots were observed.

**Supplemental_Figure_3:**

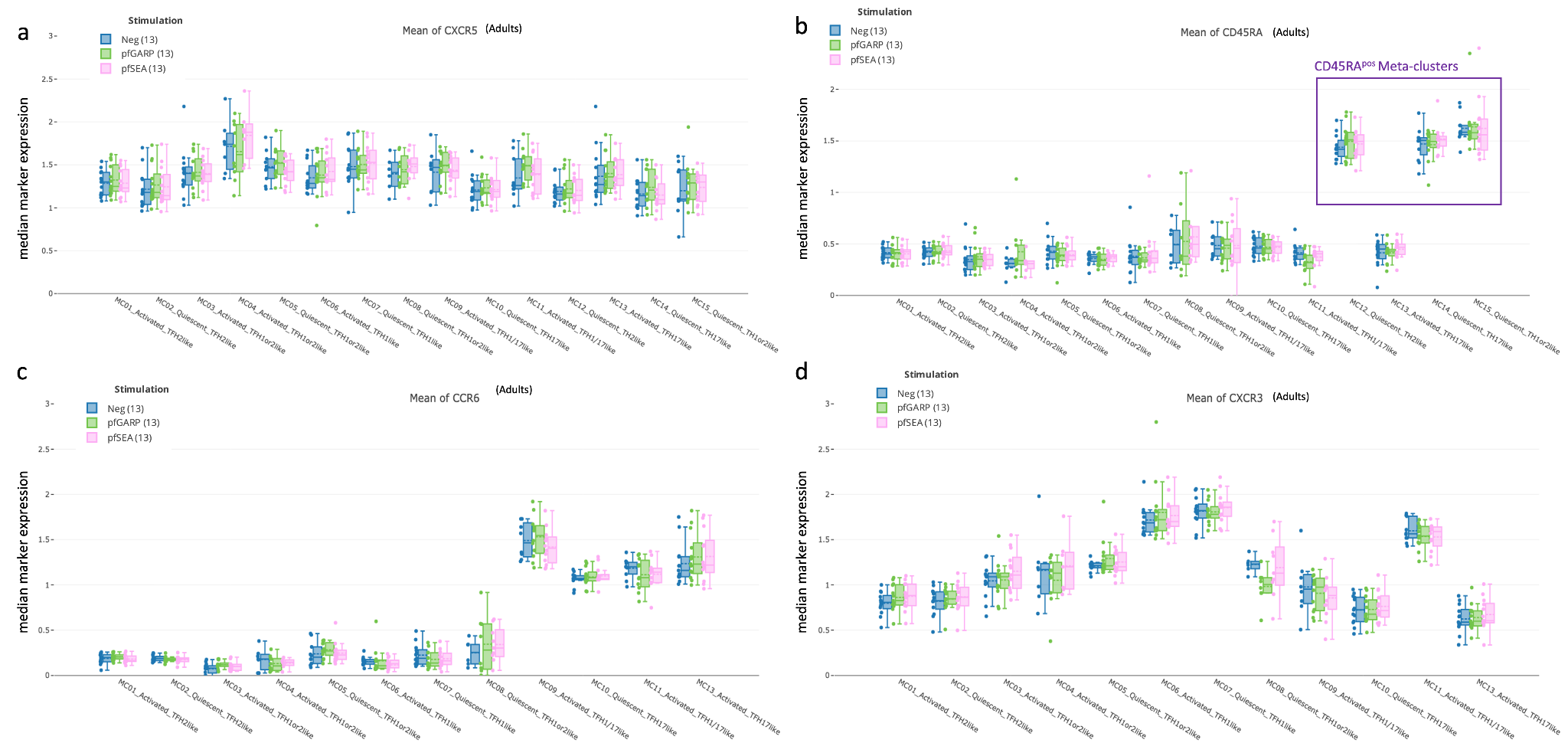

**Supplemental Figure 3: Characterization of T_FH_ subsets using CXCR5, CXCR3, CCR6, and CD45RA expression.** Bar plots showing **(a)** CXCR5, **(b)** CD45RA, **(c)** CCR6, and **(d)** CXCR3 expression across all cT_FH_-like meta-clusters from adults (n=13) under no stimulation (blue), *Pf*GARP (green) and *Pf*SEA-1A (pink) stimulations. Fifty percent of the data are within the box limits, the solid line indicates the median, the dash line the mean and the whiskers indicate the range of the remaining data with outliers being outside that range.

**Supplemental_Figure_4:**

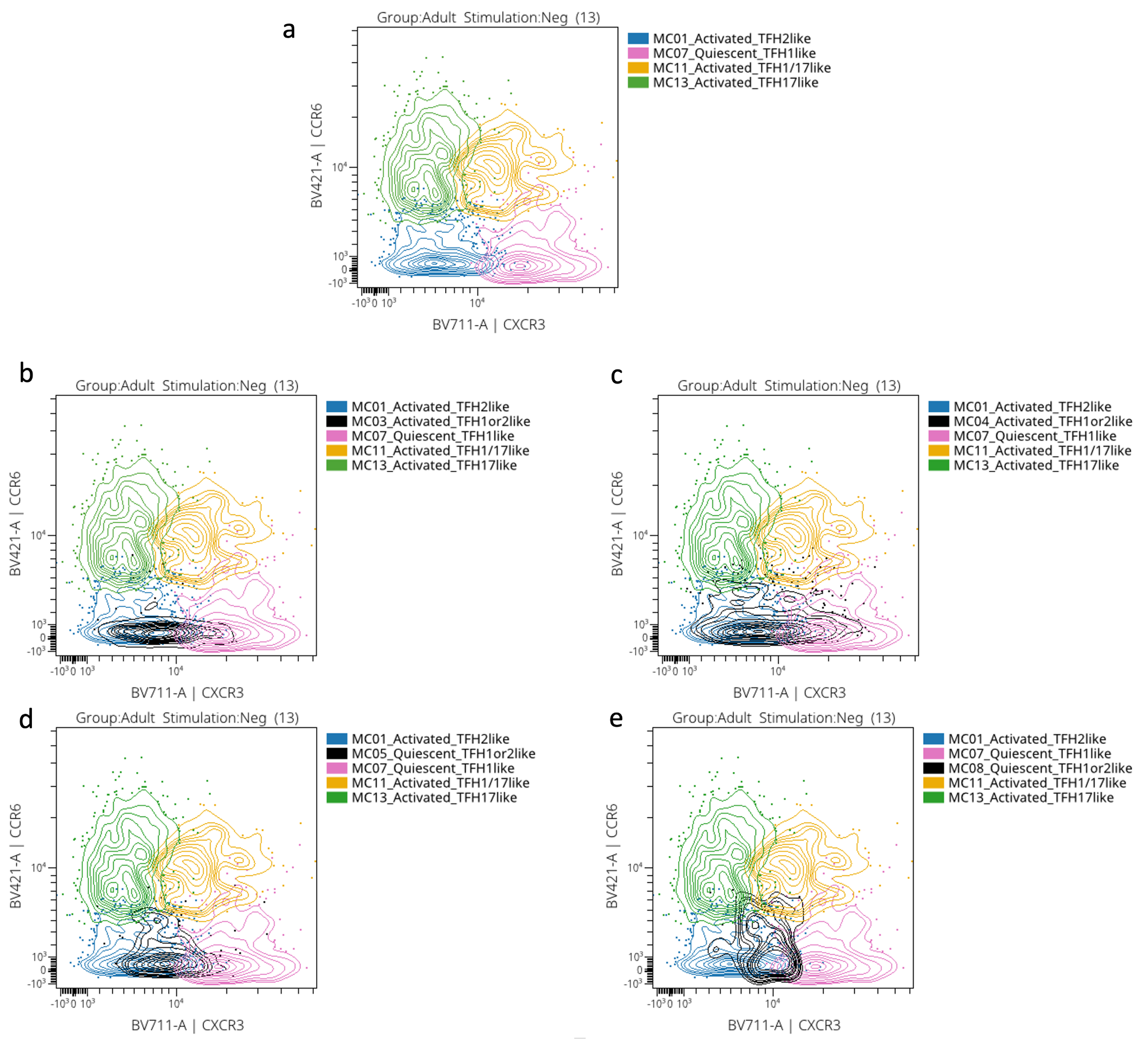

**Supplemental Figure 4: CCR6 versus CXCR3 representative cytoplots. a.** CCR6 vs CXCR3 expression from activated cT_FH_2-like MC01 (in blue), quiescent cT_FH_1-like MC07 (in pink), activated cT_FH_1/17-like MC11 (in orange) and activated cT_FH_17-like MC13 (in green). Undetermined meta-clusters was superposed to the sus-mentioned metacluster: activated cT_FH_1or2-like MC03 (in black) (**b**); activated cT_FH_1or2-like MC04 (in black) (**c**); quiescent cT_FH_1or2-like MC05 (in black) (**d**); quiescent cT_FH_1or2-like MC08 (in black) (**e**).

**Supplemental_Figure_5:**

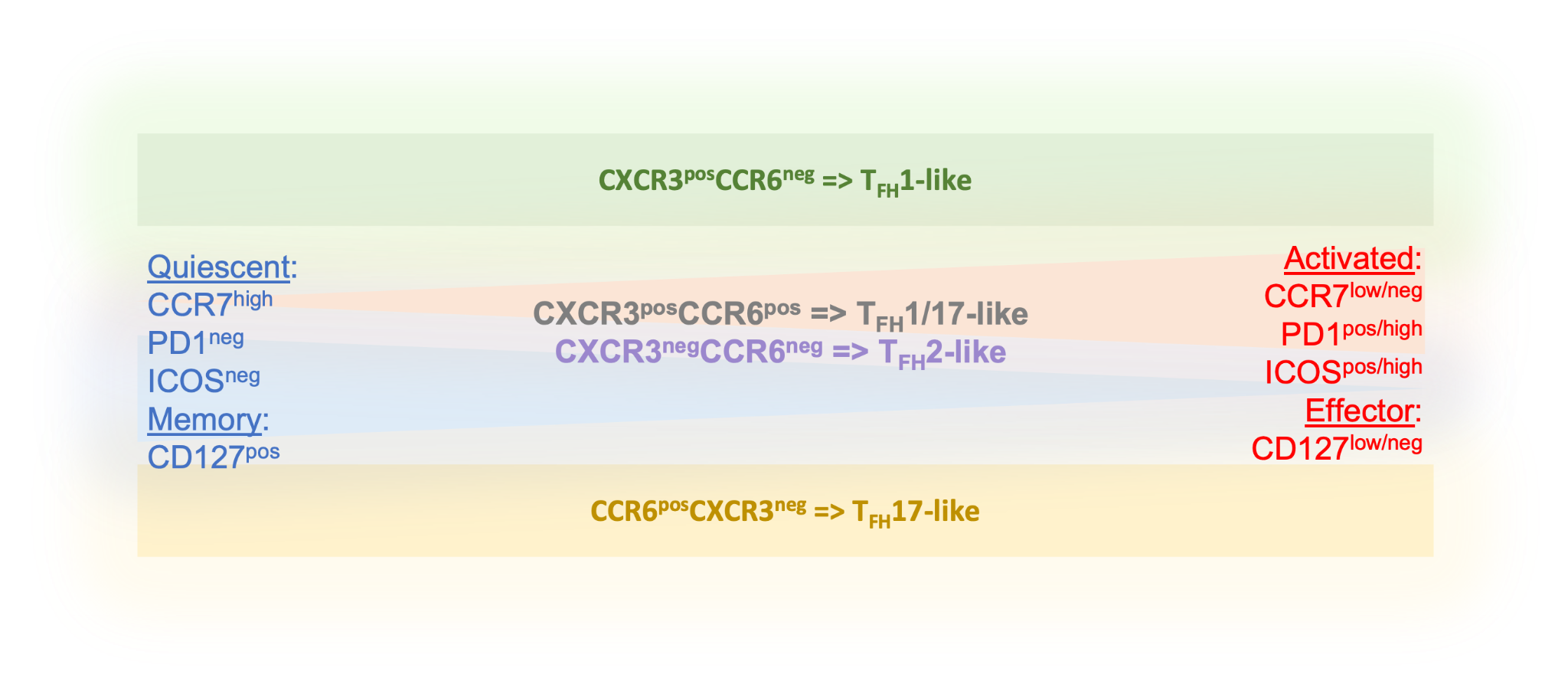

**Supplemental Figure 5: Cartoon of T_FH_ subsets and their state of activation.**

**Supplemental_Figure_6:**

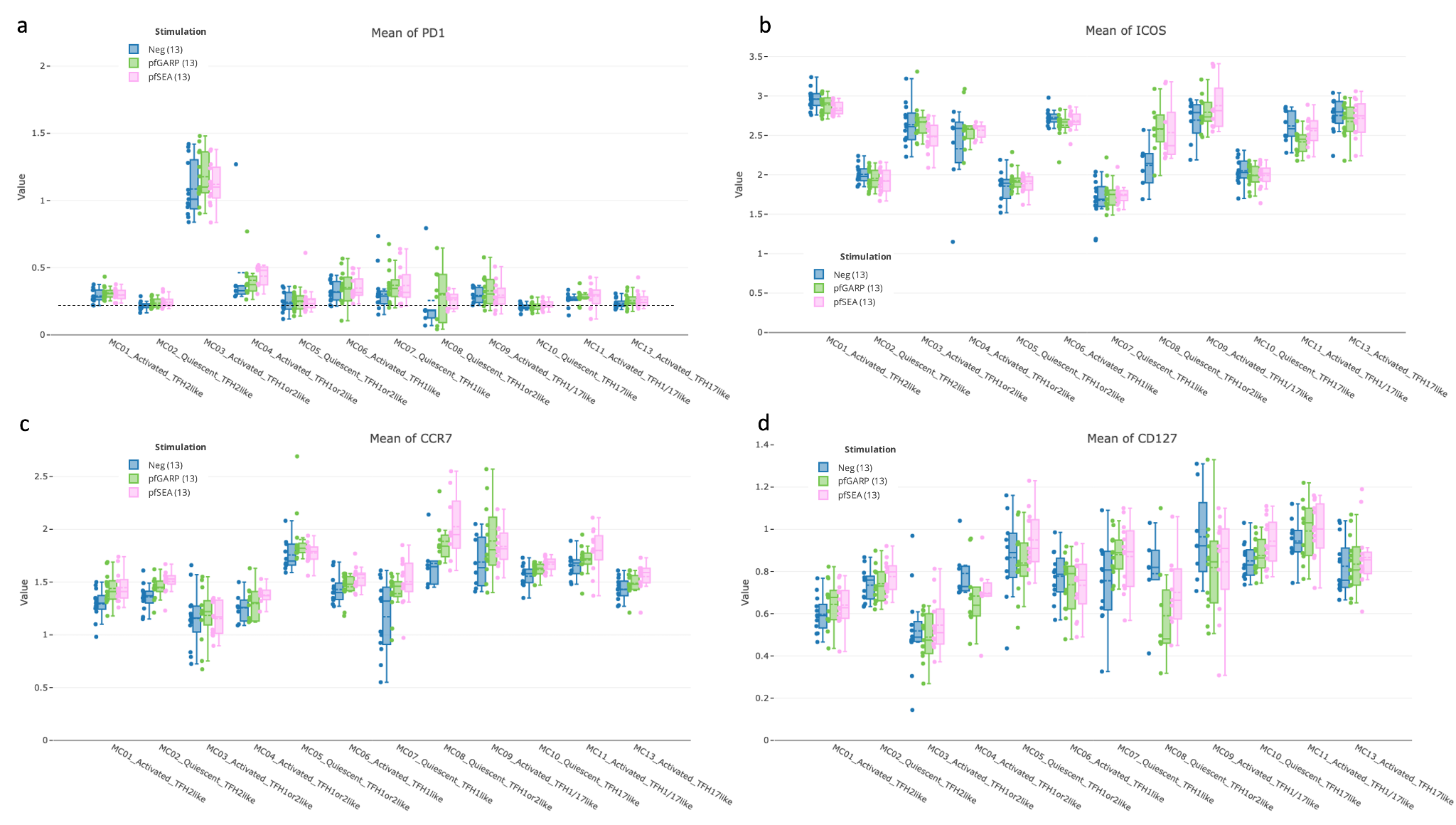

**Supplemental Figure 6: State of activation of cT_FH_ subsets from adults determined by PD1, ICOS, CCR7, and CD127.** Bar plots showing **(a)** PD1, **(b)** ICOS, **(c)** CCR7 and **(d)** CD127 expression across all cT_FH_-like meta-clusters in adults (n=13) under no stimulation (blue), *Pf*GARP (green) and *Pf*SEA-1A (pink) stimulations. Fifty percent of the data are within the box limits, the solid line indicates the median, the dash line the mean and the whiskers indicate the range of the remaining data with outliers being outside that range.

**Supplemental_Figure_7:**

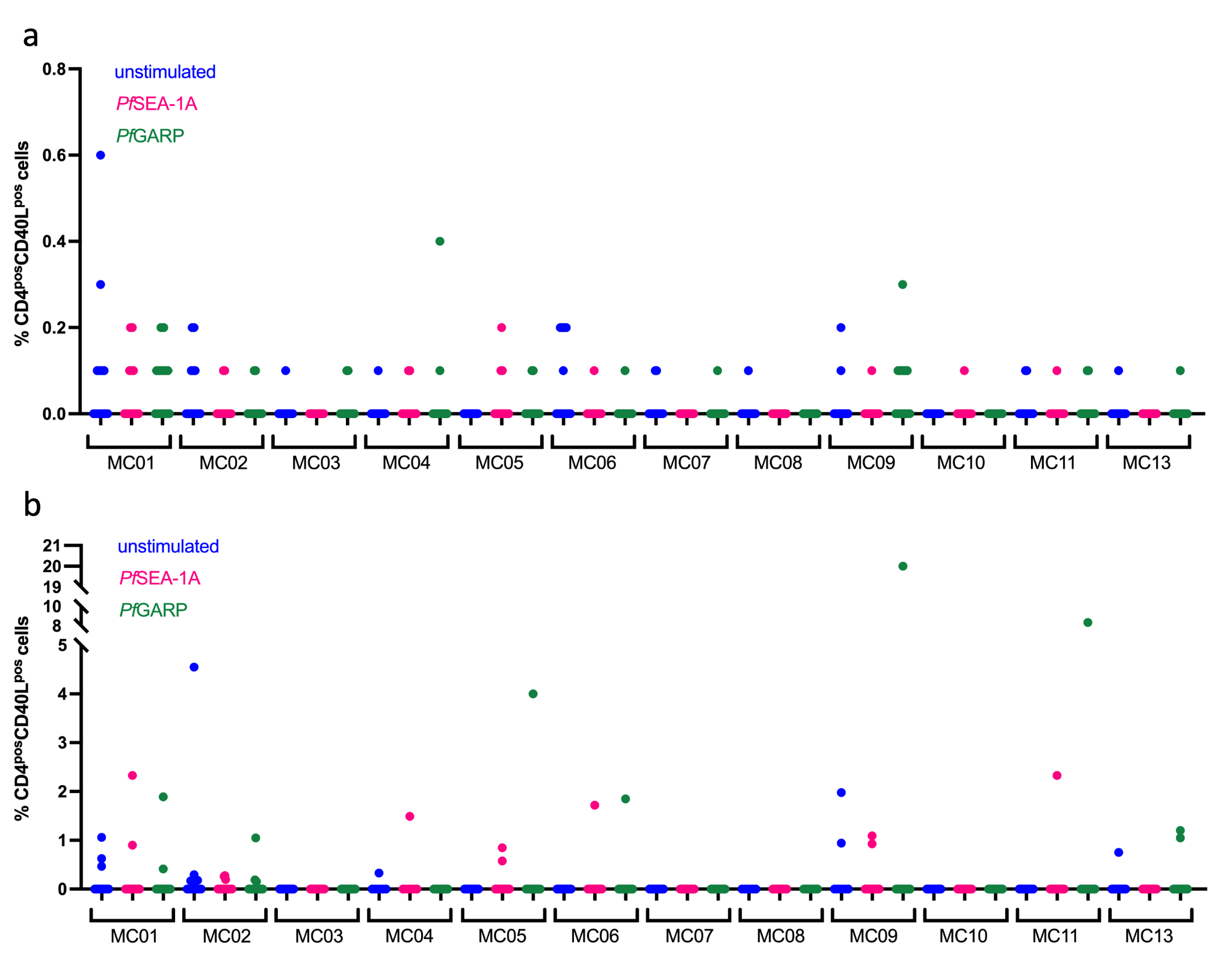

**Supplemental Figure 7: CD40L expression across stimulation and meta-clusters.** Dot plots of CD40L expression after manual gating based on unstimulated condition from children (n=13) (**a**) and adults (n=13) (**b**) cT_FH_ meta-clusters.

**Supplemental_Figure_8:**

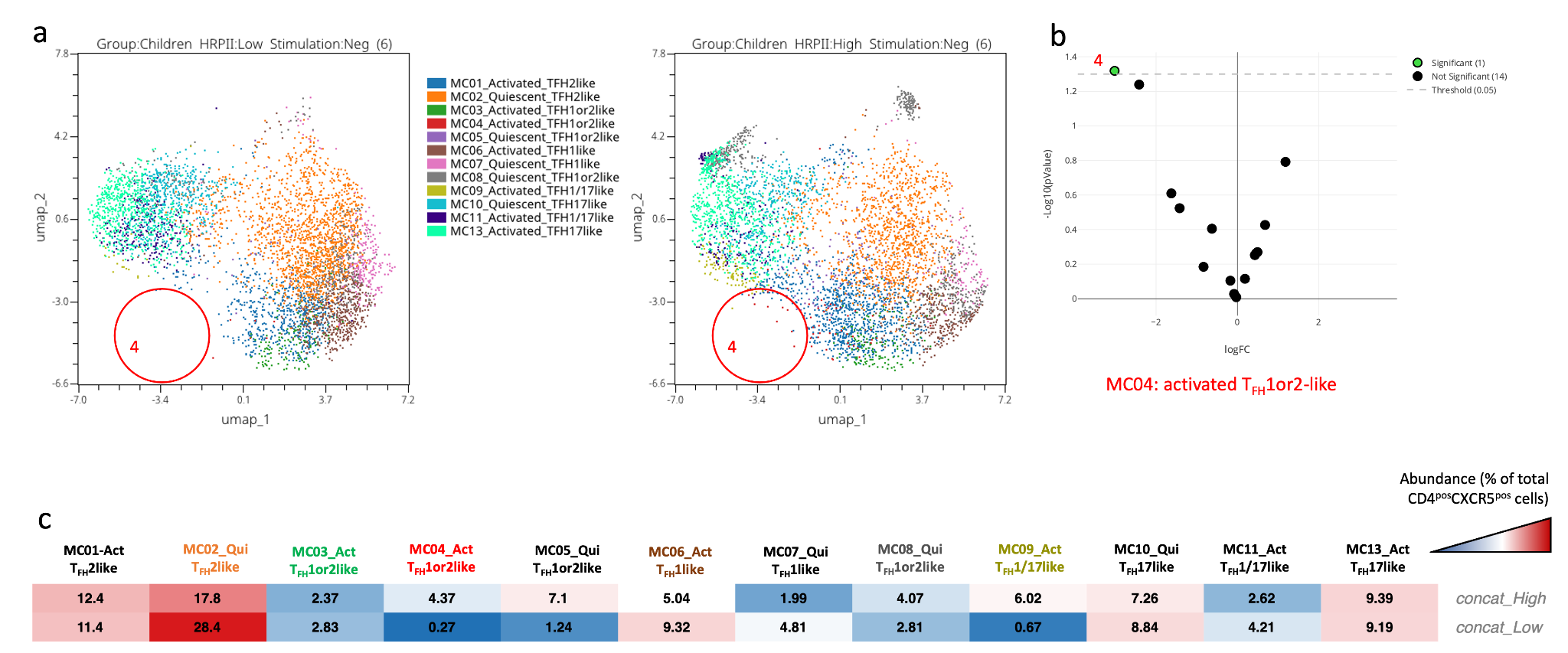

**Supplemental Figure 8: Abundance of the cT_FH_ meta-clusters within children with low or high levels of HRP2 antibodies. (a)** UMAP plot showing the clustering of the 13 different cT_FH_ meta-clusters (MC15) in children with low levels of HRP2 (on the left, n=6) and high levels of HRP2 (on the right, n=6) in the absence of stimulation. Each color represents a meta-cluster. The red circles highlight the meta-cluster showing differences in its abundance between the two groups of children. **(b)** An EdgeR statistical plot was performed to assess the abundance of the 13 meta-clusters between unstimulated PBMCs from both groups of children; the Y-axis being -log10(*p*-value) and the X-axis shows the log(FC). The green dot (MC04) is statistically significant whereas the black dots are not. **(c)** An abundance heatmap showing the distribution (in %) of each meta-cluster within the two groups of children (both concatenated unstimulated PBMCs from six participants). The color scale ranges from high abundance (red) to low abundance (blue).

**Supplemental_Figure_9:**

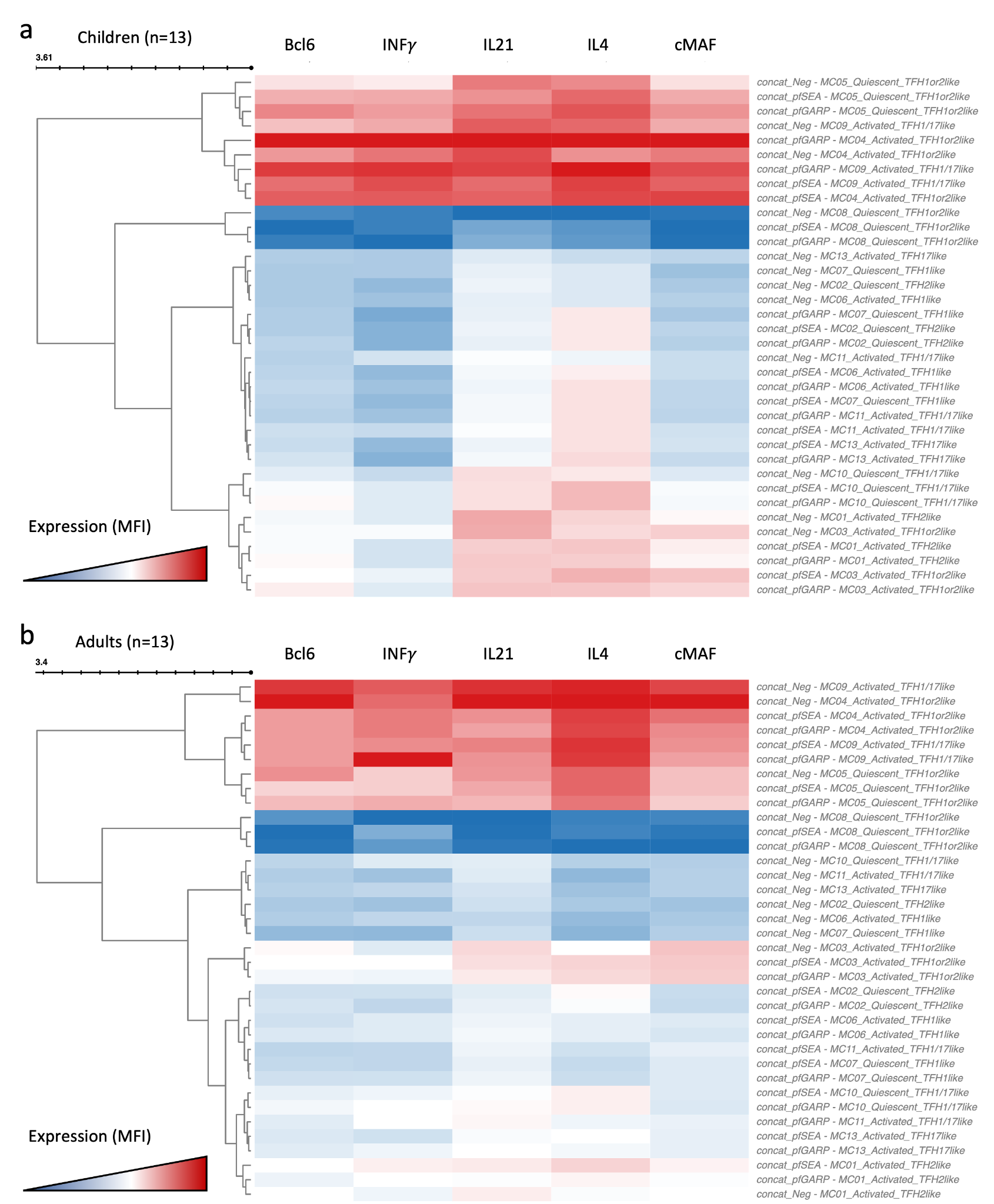

**Supplemental Figure 9: Clustered heatmap of the cytokines and transcription factors expressed from cT_FH_ meta-clusters in adults and children under the different stimulation conditions.** Heatmap of Bcl6, IFNγ, IL21, IL4, and cMAF expression from the 13 cT_FH_ meta-clusters from concatenated data from **(a)** children (n=13) and **(b)** adults (n=13), after stimulation by *Pf*SEA-1A or *Pf*GARP, or without any stimulation as indicated in the name of each row. The color scale ranges from high expression (red) to low/no expression (blue).

**Supplemental_Figure_10:**

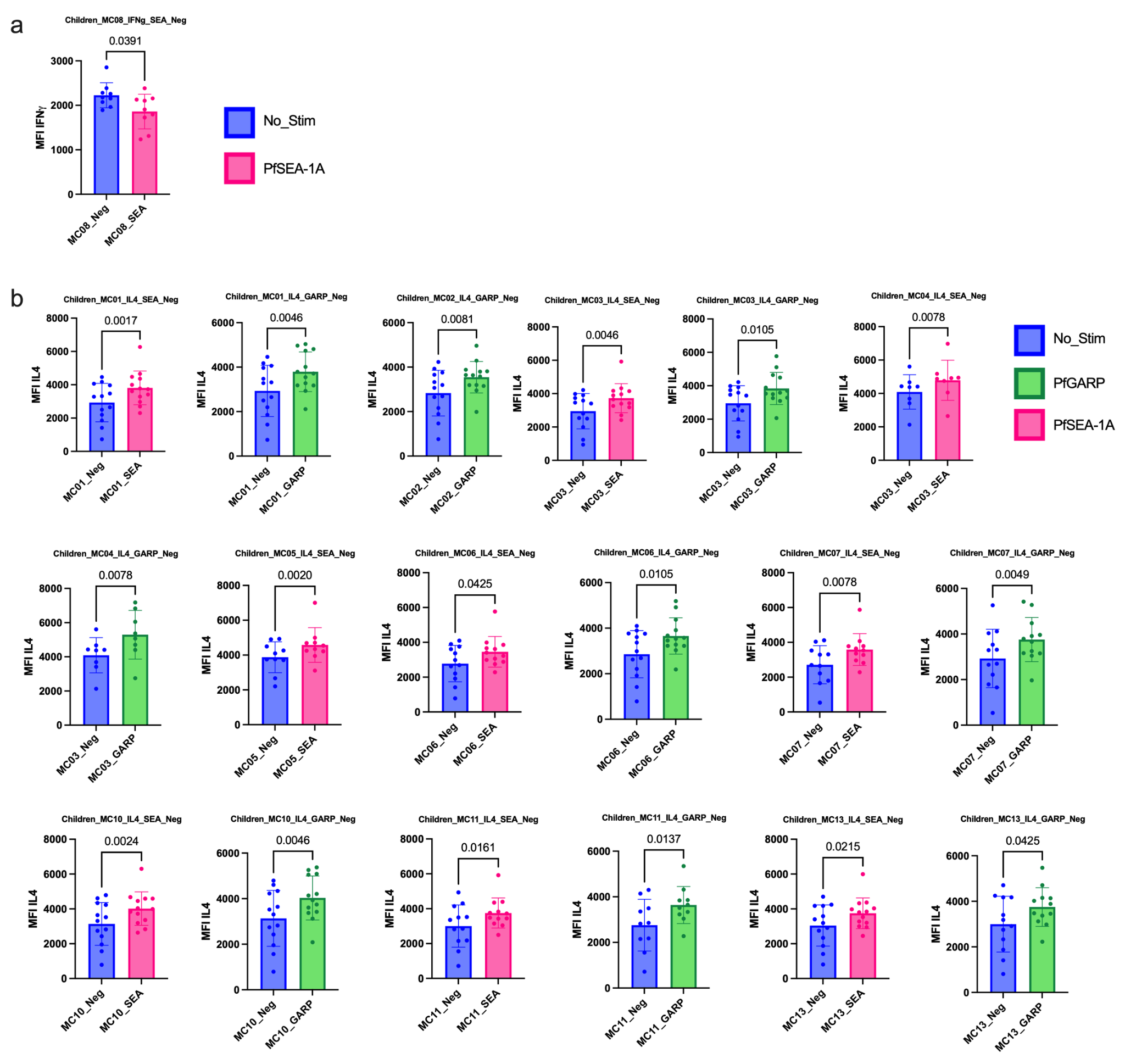

**Supplemental Figure 10: Bar plots of cytokines expressed under the different conditions of stimulation in children (n=13).** Comparison of mean fluorescent intensity (MFI) of **(a)** IFNγ, **(b)** IL4 cytokines expression after *Pf*SEA-1A (pink) or *Pf*GARP (green) stimulation or no stimulation control (blue). Bar plots indicate mean with SD. Wilcoxon-paired two-tailed t-tests were performed and *p*-values are indicated.

**Supplemental_Figure_11:**

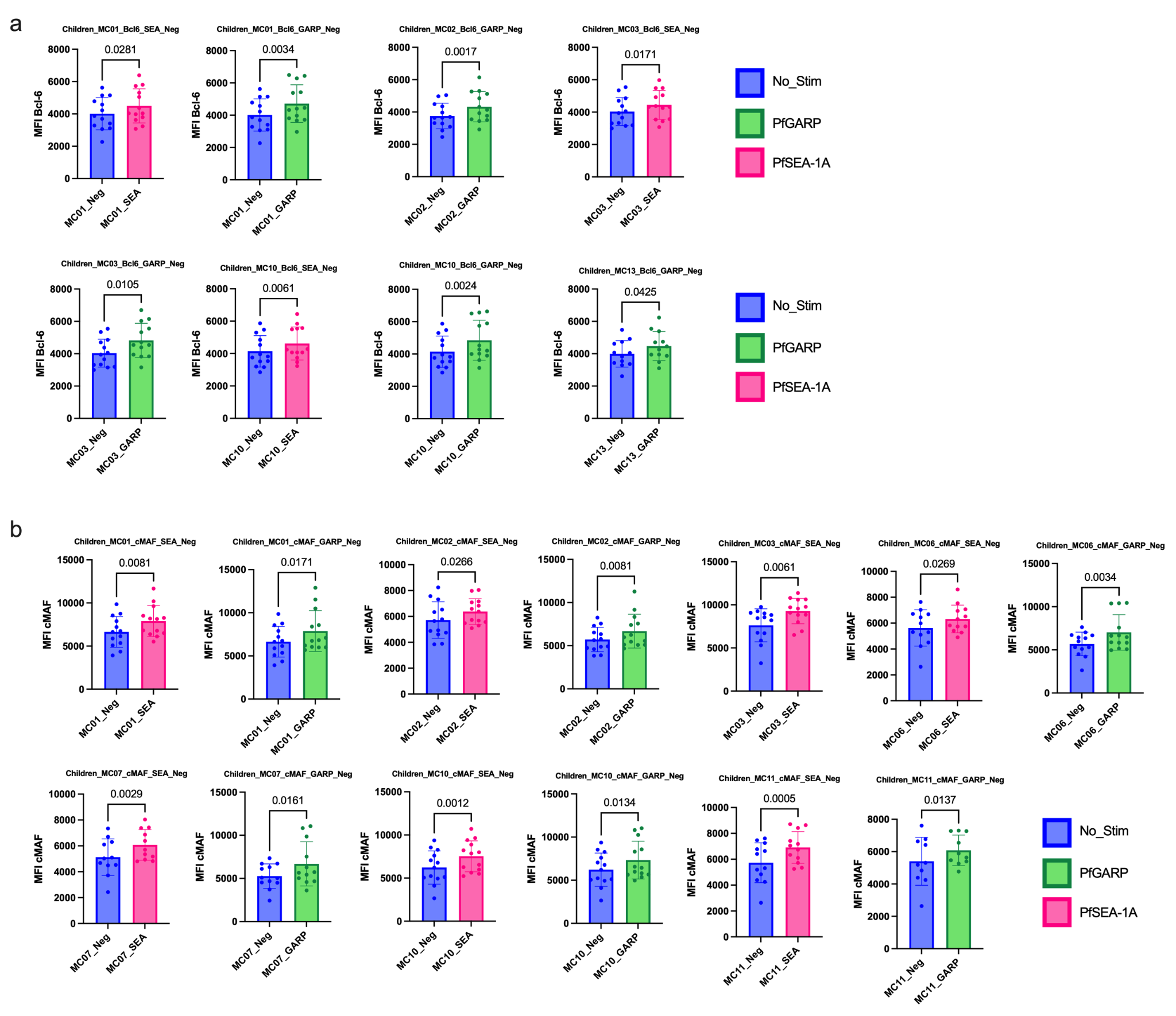

**Supplemental Figure 11: Bar plots of transcription factors expressed under the different conditions of stimulation in children (n=13).** Comparison of mean fluorescent intensity (MFI) of **(a)** Bcl6 and **(b)** cMAF expression after *Pf*SEA-1A (pink) or *Pf*GARP (green) stimulation or no stimulation control (blue). Bar plots indicate mean with SD. Wilcoxon-paired two-tailed t-tests were performed and *p*-values are indicated.

**Supplemental_Figure_12:**

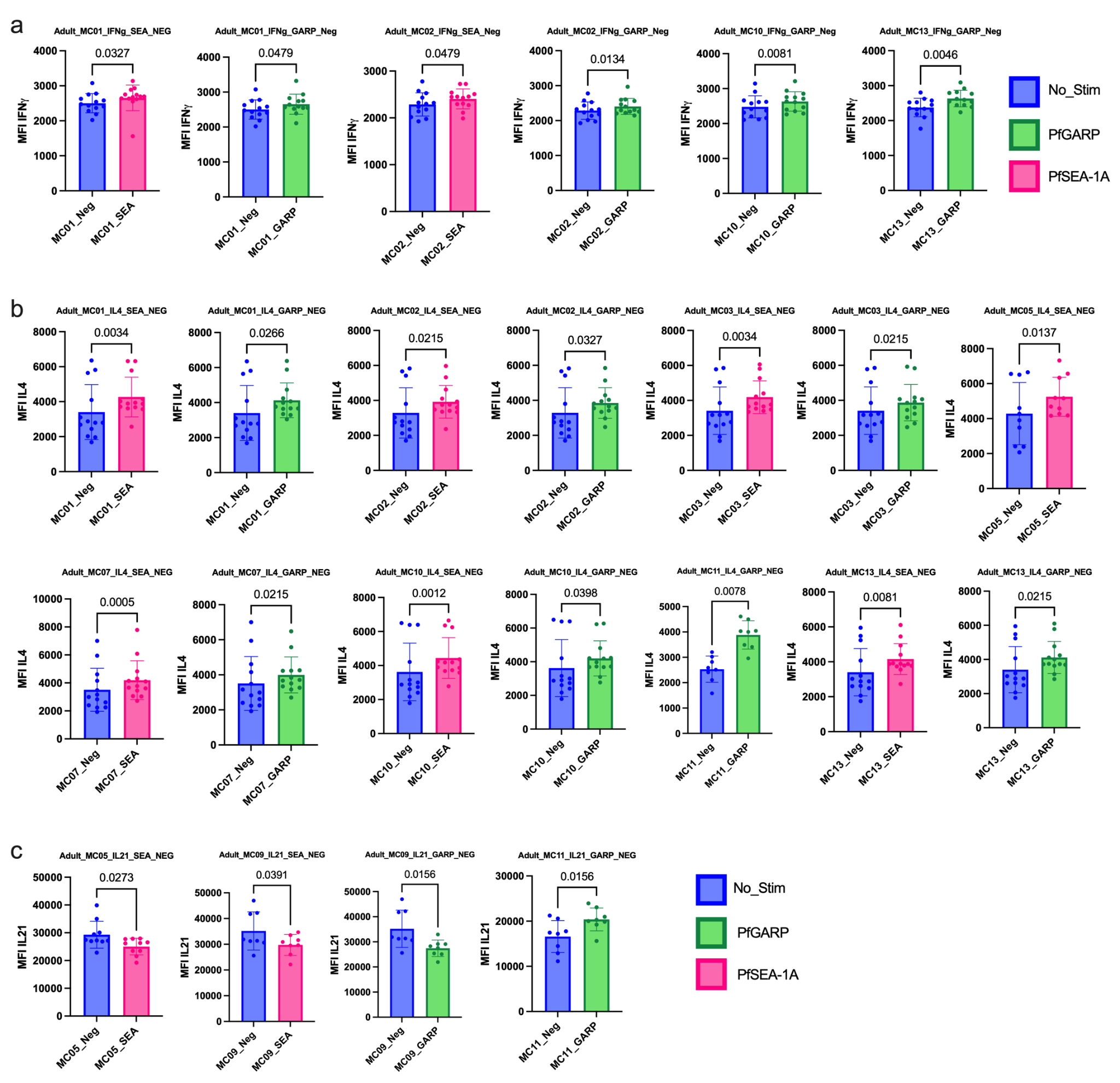

**Supplemental Figure 12: Bar plots of cytokines expressed under the different conditions of stimulation in adults (n=13).** Comparison of mean fluorescent intensity (MFI) of **(a)** IFNγ, **(b)** IL4 and **(c)** IL21 and cytokines expression after *Pf*SEA-1A (pink) or *Pf*GARP (green) stimulation or no stimulation control (blue). Bar plots indicate mean with SD. Wilcoxon-paired two-tailed t-tests were performed and *p*-values are indicated.

**Supplemental_Figure_13:**

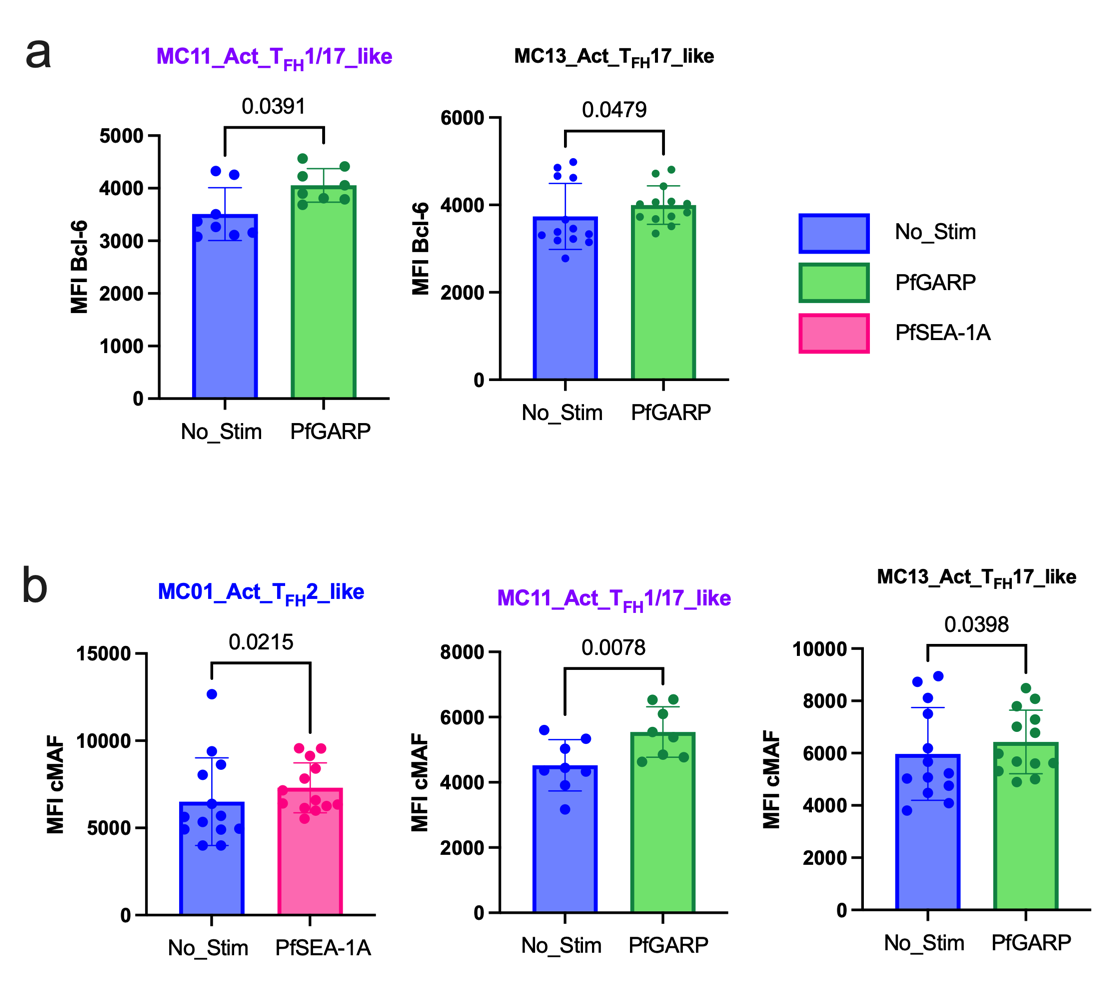

**Supplemental Figure 13: Bar plots of transcription factors expressed under the different conditions of stimulation in adults (n=13).** Comparison of mean fluorescent intensity (MFI) of **(a)** Bcl6 and **(b)** cMAF expression after *Pf*SEA-1A (pink) or *Pf*GARP (green) stimulation or no stimulation control (blue). Bar plots indicate mean with SD. Wilcoxon-paired two-tailed t-tests were performed and *p*-values are indicated.

**Supplemental_Figure_14:**

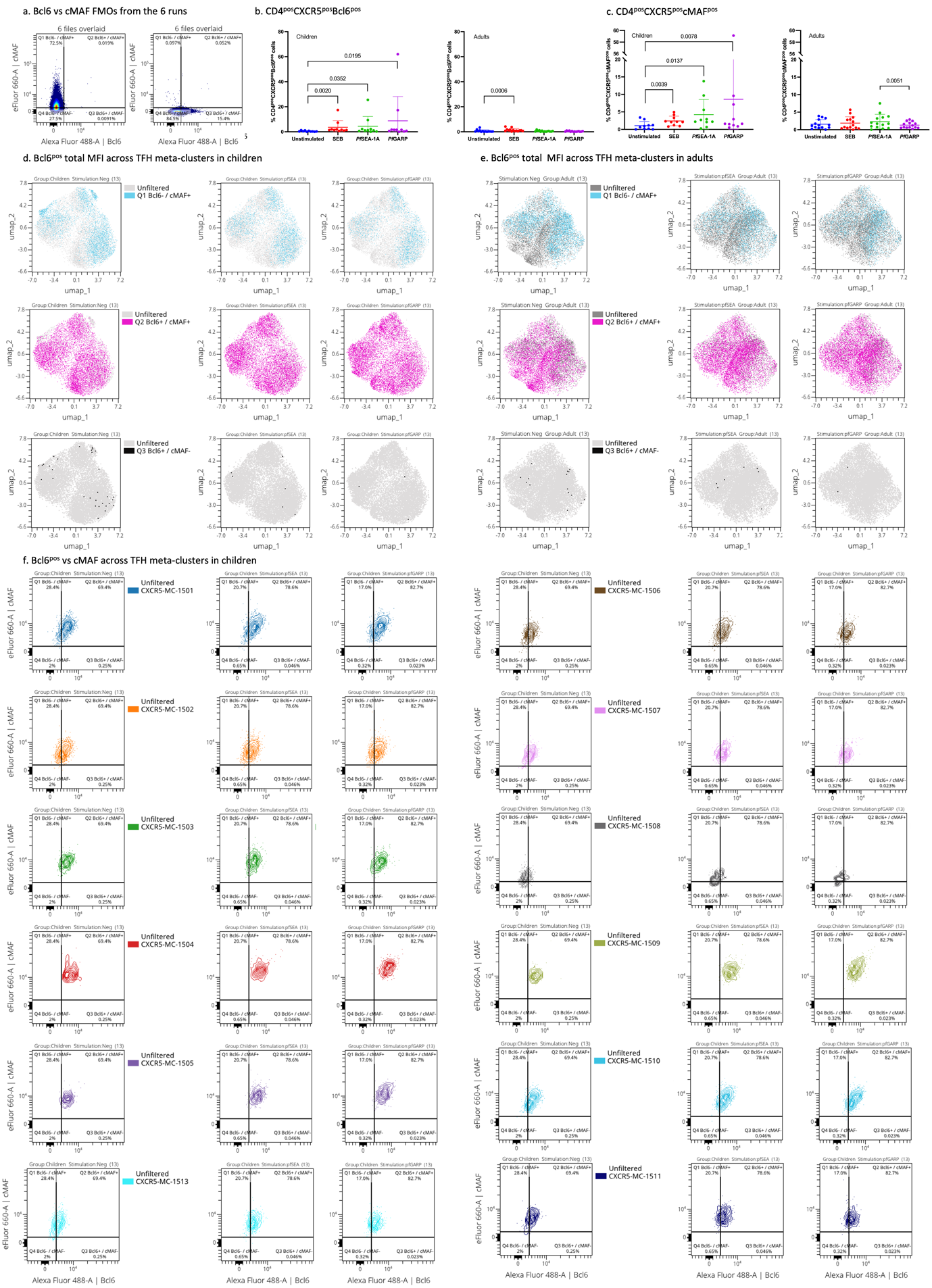

**Supplemental Figure 14: Manually gated transcription factors expression upon stimulation. A)** Bcl6 vs cMAF cytoplots from the 6 runs overlaid FMOs. **B)** Percentage of CD4^pos^CXCR5^pos^Bcl6^pos^ cells across stimulation in children (left) and adults (right). Bar plots indicate mean with SD. Wilcoxon-paired two-tailed t-tests were performed and *p*-values are indicated. **C)** Percentage of CD4^pos^CXCR5^pos^cMAF^pos^ cells across stimulation in children (left) and adults (right). Bar plots indicate mean with SD. Wilcoxon-paired two-tailed t-tests were performed and *p*-values are indicated. Manually gated positive Bcl6 cells overlaid on the UMAP of the CD4^pos^CXCR5^pos^CD25^neg^ cells across stimulation in children **(D)** and adults **(E)**. **F)** Contour plots with outliers of cMAF vs Bcl6 expression upon stimulation for each meta-cluster.

**Supplemental_Figure_15:**

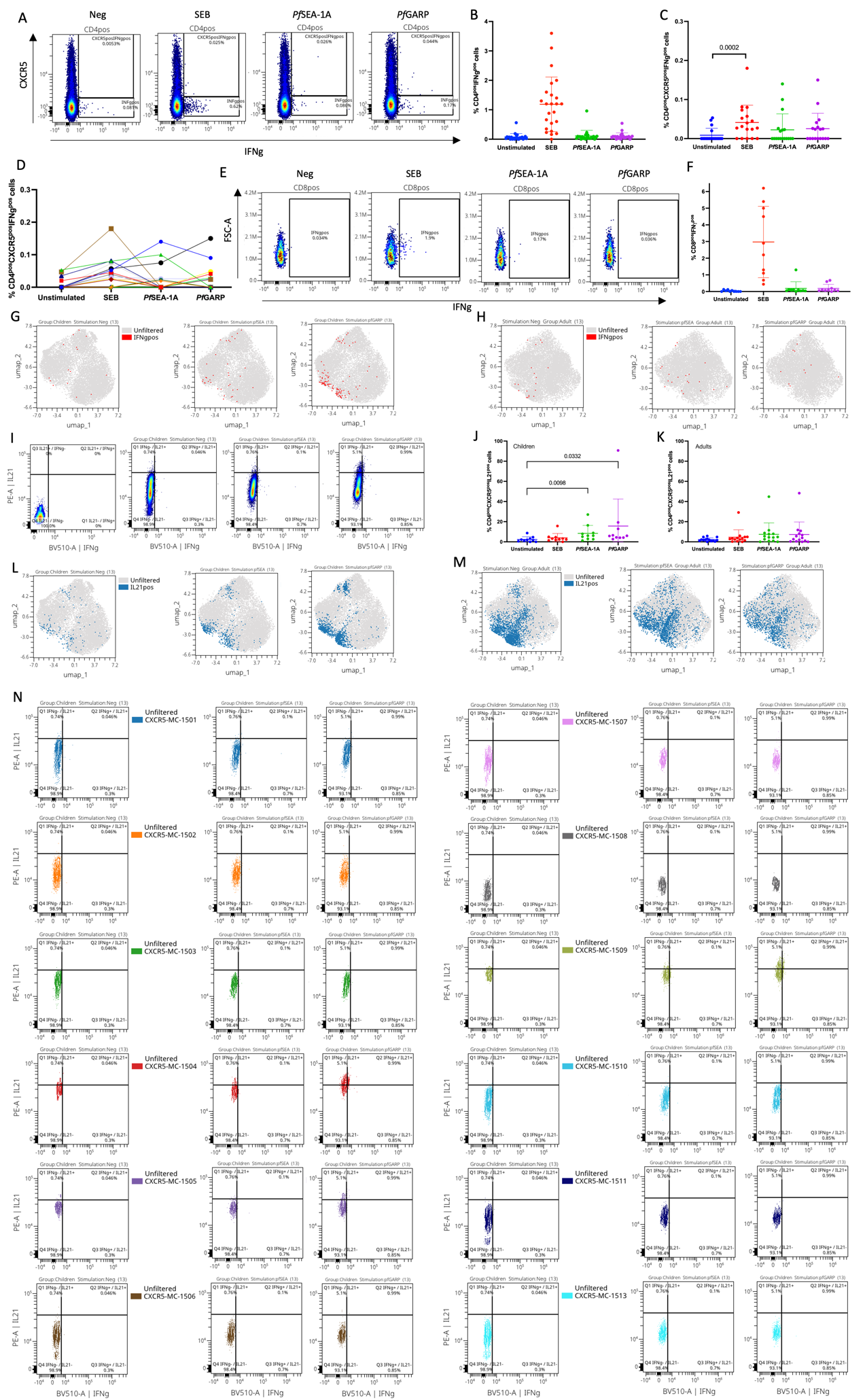

**Supplemental Figure 15: Manually gated IFNγ and IL21 expression upon stimulation. A)** Cytoplot of IFNγ vs CXCR5 expression from CD4 T-cells upon stimulation. **B)** Percentage of CD4^pos^IFNγ^pos^ cells across stimulation. **C)** Percentage of CD4^pos^CXCR5^pos^IFNγ^pos^ cells across stimulation. **D)** Spaghetti plot of the CD4^pos^CXCR5^pos^IFNγ^pos^ cells across stimulation, each color represents a participant. **E)** Cytoplots of IFNγ vs FSC-A expression from CD89 T-cells upon stimulation. **F)** Percentage of CD8^pos^IFNγ^pos^ cells across stimulation. Manually gated positive IFNγ cells overlaid on the UMAP of the CD4^pos^CXCR5^pos^CD25^neg^ cells across stimulation in children **(G)** and adults **(H)**. **I)** Cytoplots of IFNγ vs IL21 expression from 6 overlaid FMOs and CD4^pos^CXCR5^pos^ T-cells upon stimulation. **J)** Percentage of CD4^pos^IL21^pos^ cells across stimulation in children. **K)** Percentage of CD4^pos^IL21^pos^ cells across stimulation in adults. Manually gated positive IL21 cells overlaid on the UMAP of the CD4^pos^CXCR5^pos^CD25^neg^ cells across stimulation in children **(L)** and adults **(M)**. **N)** Contour plots with outliers of IFNγ vs IL21 expression upon stimulation for each meta-cluster. Bar plots indicate mean with SD. Wilcoxon-paired two-tailed t-tests were performed and *p*-values are indicated.

**Supplemental_Figure_16:**

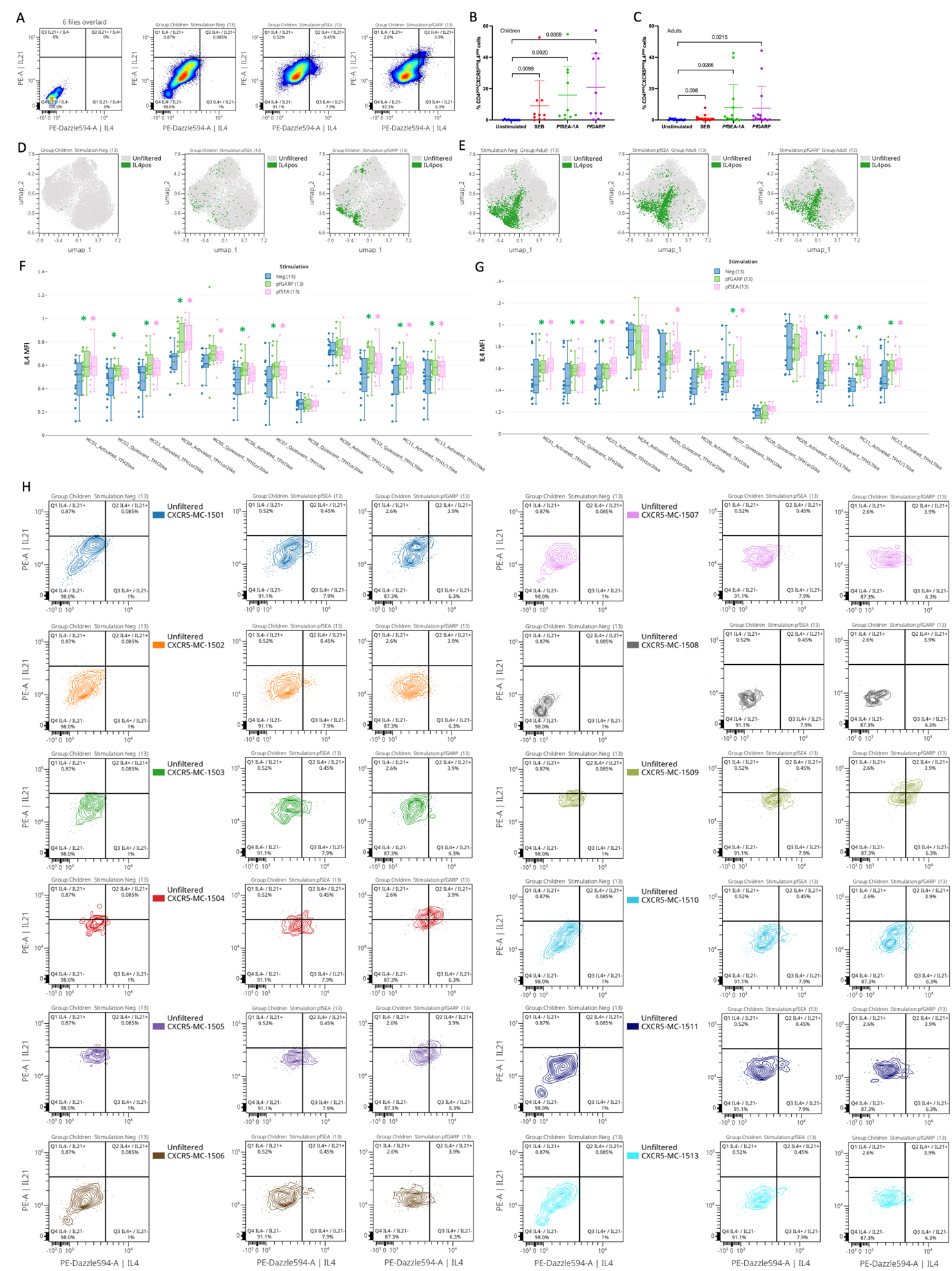

**Supplemental Figure 16: Manually gated IL4 and IL21 expression upon stimulation. A)** Cytoplot of IL4 vs IL21 expression from 6 overlaid FMO and CD4^pos^CXCR5^pos^ T-cells upon stimulation. Bar plots indicate mean with SD. Wilcoxon-paired two-tailed t-tests were performed and *p*-values are indicated. Percentage of CD4^pos^CXCR5^pos^IL4^pos^ cells across stimulation in children **(B)** and adults **(C)**. Manually gated positive IL4 cells overlaid on the UMAP of the CD4^pos^CXCR5^pos^CD25^neg^ cells across stimulation in children **(D)** and adults **(E)**.

IL4 MFI for all meta-clusters represented in bar plots across stimulation in children **(F)** and adults **(G)**, negative control in blue, PfSEA-1A in pink and PfGARP in green. Wilcoxon-paired two-tailed t-tests were performed, and significant *p*-values are indicated with * equal *p*<0.05. **H)** Contour plots with outliers of L4 vs IL21 expression upon stimulation for each meta-cluster.
